# Chikungunya Virus Release is Reduced by TIM-1 Receptors Through Binding of Envelope Phosphatidylserine

**DOI:** 10.1101/2024.01.25.577233

**Authors:** Judith M. Reyes Ballista, Ashley J. Hoover, Joseph T. Noble, Marissa D. Acciani, Kerri L. Miazgowicz, Sarah A. Harrison, Grace Andrea L. Tabscott, Avery Duncan, Don N. Barnes, Ariana R. Jimenez, Melinda A. Brindley

## Abstract

T-cell immunoglobin and mucin domain protein-1 (TIM-1) mediates entry of Chikungunya virus (CHIKV) into some mammalian cells through the interaction with envelope phospholipids. While this interaction enhances entry, TIM has been shown to tether newly formed HIV and Ebola virus particles, limiting their efficient release. In this study, we investigate the ability of surface receptors such as TIM-1 to sequester newly budded virions on the surface of infected cells. We established a luminescence reporter system to produce Chikungunya viral particles that integrate nano-luciferase and easily quantify viral particles. We found that TIM-1 on the surface of host cells significantly reduced CHIKV release efficiency in comparison to other entry factors. Removal of cell surface TIM-1 through direct cellular knock-out or altering the cellular lipid distribution enhanced CHIKV release. Over the course of infection, CHIKV was able to counteract the tethering effect by gradually decreasing the surface levels of TIM-1 in a process that appears to be mediated by the nonstructural protein 2. This study highlights the importance of phosphatidylserine receptors in mediating not only the entry of CHIKV but also its release and could aid in developing cell lines capable of enhanced vaccine production.

**Importance:** Chikungunya virus (CHIKV) is an enveloped alphavirus transmitted by the bites of infectious mosquitoes. Infection with CHIKV results in the development of fever, joint pain, and arthralgia that can become chronic and last for months after infection. Prevention of this disease is still highly focused on vector control strategies. In December 2023, a new live attenuated vaccine against CHIKV was approved by the FDA. We aimed to study the cellular factors involved in CHIKV egress, to better understand CHIKV’s ability to efficiently infect and spread among a wide variety of cell lines. We found that TIM-1 receptors can significantly abrogate CHIKV’s ability to efficiently exit infected cells. This information can be beneficial for maximizing viral particle production in laboratory settings and during vaccine manufacturing.

## INTRODUCTION

Chikungunya virus (CHIKV) is an enveloped positive-sense RNA virus from the *Togaviridae* family. Within the alphavirus genus, CHIKV causes the most human infections and is transmitted by the bites of infectious *Aedes aegypti* and *Aedes albopictus* mosquitoes (1). Chikungunya disease presents with fever, joint pain, stiffness, and arthralgia with some patients experiencing severe joint pain for months after infection. During outbreaks, efforts to slow transmission and spread focused on decreasing the mosquito vector populations. In December 2023, the FDA approved a new live attenuated vaccine for the prevention of CHIKV which will hopefully aid in slowing future outbreaks. We still lack antivirals to treat CHIKV infection, therefore, identifying potential targets of CHIKV replication cycle may provide new targets for the development of new therapeutics.

Chikungunya virus encodes four non-structural proteins (nsP1-nsP4) and five structural and accessory proteins (C, E3, E2, 6k, TF, and E1). While the non-structural proteins are responsible for transcription and genome replication, the structural proteins assemble to form particles. CHIKV particles are composed of a nucleocapsid core comprised of the RNA genome and capsid proteins, surrounded by a lipid envelope studded with glycoproteins (2). Structural studies observed the envelope E1-E2 spikes organized in a hexagonal lattice at the plasma membrane, the site of virus budding (3). As the nucleocapsid buds through the plasma membrane, both the capsid and glycoproteins are arranged in icosahedral shells (T=4) (3). Assembly of CHIKV is driven by the interaction of capsid protein and the cytoplasmic tail of the E2 protein (4–7). However, recent studies suggest that accessory proteins 6k and TF may also facilitate efficient exit of Sindbis virus, a closely related alphavirus (8).

While capsid and E2 interactions initiate CHIKV particle budding, subsequent events involving additional cellular proteins may be required for efficient release. Many enveloped viruses utilize the cellular endosomal sorting complexes required for transport (ESCRT) proteins to complete the final membrane scission (9). Once newly formed particles are separated from the plasma membrane, the particles need to escape from the infected cell to perpetuate infection. The interferon-induced transmembrane protein, tetherin/BST-2, is a cellular surface protein that can inhibit the release of enveloped viruses including human immunodeficiency virus (HIV), Ebola, and CHIKV (10–13). Although previous studies have shed light on the mechanisms of alphavirus budding, our knowledge of cellular factors that can alter particle release is limited.

Phosphatidylserine (PS) in the lipid envelope of viral particles influences multiple steps of the viral replication cycle (14). PS is an anionic phospholipid typically found on the inner leaflet of the plasma membrane (15). Apoptotic cells move PS to the outer leaflet which serves as a marker for cell clearance (16). This process is mediated by flippases, which translocate PS from the outer leaflet to the inner leaflet of the plasma membrane, and scramblases which non-specifically shuffle PS between the leaflets. Viral envelopes rich in PS can enter cells via apoptotic mimicry, where outer leaflet PS in the viral envelope attaches to PS receptors on the surface of the host cells. Our group and others showed that CHIKV entry in certain cell lines (*i.e.,* Vero cells) is mainly mediated through attachment to PS receptors, such as T cell immunoglobulin mucin domain-1 (TIM-1) and receptor tyrosine kinase AXL (AXL) (17–19). Increased levels of PS in CHIKV’s outer leaflet enhanced the specific infectivity of particles into Vero cells (17). While PS receptors can aid in virus entry, they can also modulate immune responses and reduce virion release. For example, PS receptors can prevent the efficient release of HIV and Japanese Encephalitis virus (JEV) by attaching to the viral envelope PS in newly budded particles (20, 21). To our knowledge, no previous studies have noted the role of PS receptors in the viral particle release of alphaviruses.

In this study, we aimed to understand the role of PS receptors in the release of CHIKV particles. To facilitate viral quantification, we utilized CHIKV tagged with a nano-luciferase directly integrated into viral particles. We found that cells lacking TIM-1 receptors released more CHIKV particles in comparison to their wildtype counterparts which sequestered particles through TIM-1 interaction. Likewise, cells producing exogenous TIM-1 released fewer particles. The change in particle release was directly attributed to the PS binding domain in TIM-1. Chikungunya infection counteracts this effect by reducing surface TIM levels as infection proceeds. Cells producing nsP2 displayed reduced PS binding activity, suggesting it plays a role in counteracting TIM-1 in particle release. This study highlights an additional role of PS receptors in the CHIKV replication cycle.

## RESULTS

### Chikungunya virus exhibits an increased release efficiency in Vero cells lacking PS receptors

We previously demonstrated that CHIKV entry in Vero cells is facilitated by PS receptors (17). While entry into Vero cells lacking both TIM-1 and AXL (VeroΔTIM/AXL) was inefficient, the amount of virus produced from the cells was higher than expected. For example, when Vero and VeroΔTIM/AXL cells are infected with an equivalent amount of CHIKV particles, viral protein was poorly detected in VeroΔTIM/AXL cell lysates and more readily detected in the supernatant containing released viral particles. (Figure 1A). If VeroΔTIM/AXL cells are infected with ten times more virions, Vero and VeroΔTIM/AXL cells display comparable levels of E1 protein (Figure 1A). We observed more CHIKV E1 protein in the supernatant from VeroΔTIM/AXL than in parental cells and the release efficiency (ratio of protein levels in the supernatants over cell lysates) was increased 4-fold in ΔTIM/AXL cells (Figure 1A). This data suggests that while PS receptors mediate CHIKV entry into Vero cells, they can also decrease particle release.

**Figure 1.**
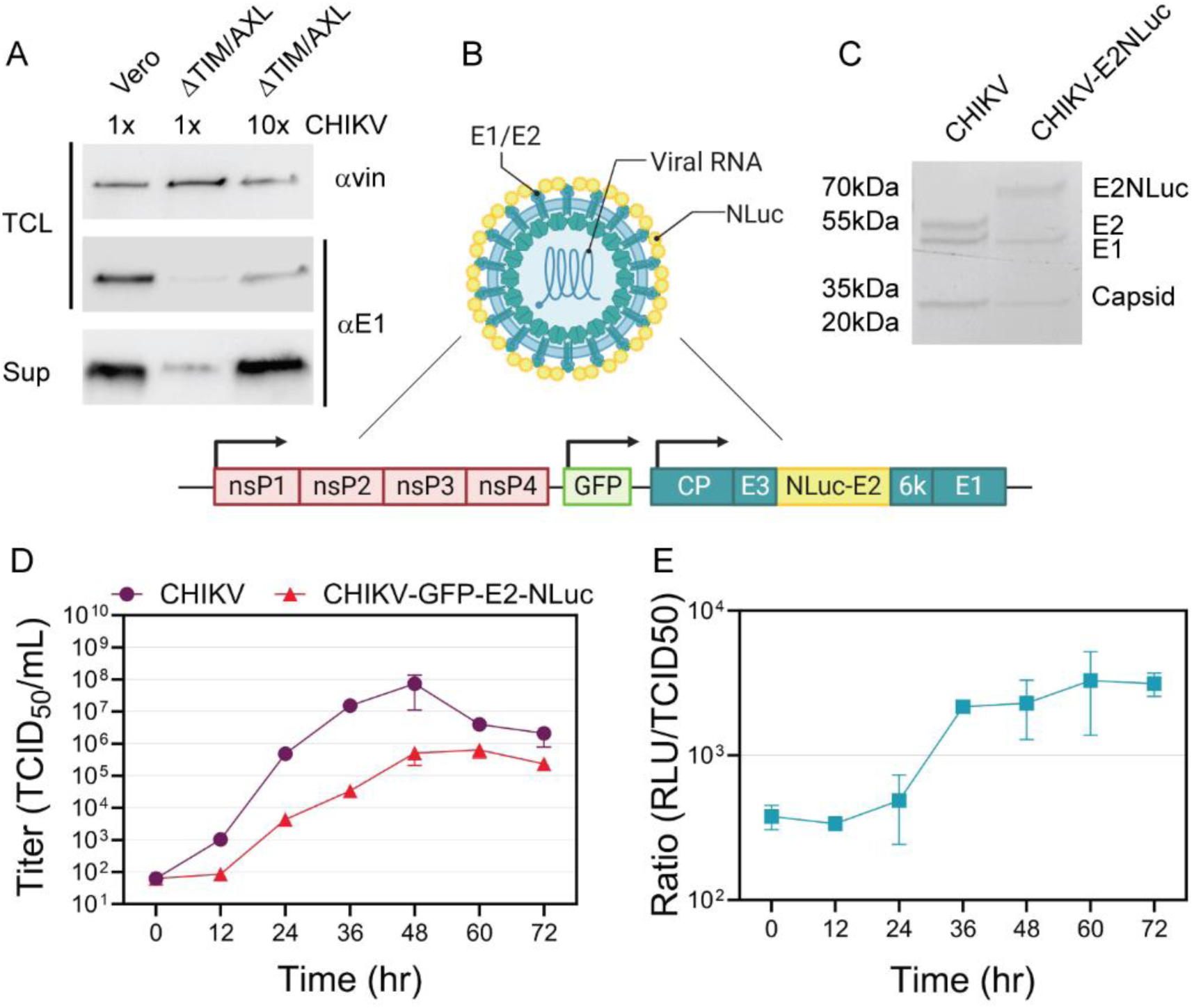
Nano luciferase tag serves as a measure for quantification of CHIKV viral particles. **(A)** Immunoblot analysis of total lysates and supernatants harvested from CHIKV-infected Vero and VeroΔTIM/AXL cells. Cells were infected with CHIKV-GFP at 0.5 (1x) or 5 (10x) MOI. Total cell lysates and purified supernatants were probed against CHIKV E1 or vinculin as a control. **(B)** Diagram of CHIKV-GFP-E2-NLuc virus genome used for release efficiency assays. Nano luciferase (NLuc) was inserted at the N-terminus of E2. Created in BioRender.com **(C)** Vero cells were infected with either CHIKV-GFP or CHIKV-GFP-E2-NLuc. Supernatants were purified through ultracentrifugation and analyzed using a stain-free gel. **(D)** Multi-step replication curve of CHIKV and CHIKV-GFP-E2-NLuc in Vero cells (0.01 MOI) was harvested at each indicated time point. **(E)** Ratio between TCID50U/mL and Relative Luminescence Units (RLU) from samples harvested in the multi-step replication curve of cells infected with CHIKV-GFP-E2-NLuc. Data represents the mean ±SEM from at least three independent trials.

To more readily quantify CHIKV viral release efficiency, we cloned nano-luciferase (NLuc) to the N-terminus of the E2 glycoprotein as previously described (Figure 1B) (22). The recombinant virus contains NLuc in the virion, therefore viral particles can be readily quantified using a standard luminescence assay. Purified CHIKV particles displayed three proteins (capsid, E1, and E2) and showed an increase of ∼20kDa in the E2 protein, corresponding to the NLuc enzyme (Figure 1C). Each particle theoretically incorporates 240 NLuc attached to each E2 molecule in the particle. Similar ratios of capsid:E1:E2 were observed in both the parental and tagged viruses suggesting NLuc incorporation did not impede or alter particle formation (Figure 1C). While tagging the virus reduced CHIKV titers, it did not significantly alter the replication kinetics. Both tagged and untagged virus titers peaked around 48 hours after infection (Figure 1D). When comparing the NLuc levels to infectious titers over time (Figure 1E), we observed a consistent ratio of ∼2700±500 RLU/TCID50 U once infection was established (36hpi onward), suggesting consistent luciferase activity is associated with infectious virions.

To further investigate the role of PS receptors on CHIKV viral release, we infected Vero cells knocked out for TIM-1, AXL, or both TIM-1 and AXL with CHIKV-GFP-E2-NLuc and calculated release efficiency by comparing the luciferase activity present in the supernatant to the cell lysate levels. We observed that CHIKV particles were 2-3 times more efficiently released in VeroΔTIM and VeroΔTIM/AXL cells than in the parental cell line (Figure 2A). To ensure that this was not an artifact from the entry defect observed in cells lacking TIM-1 and AXL (Figure 2B), we repeated the experiment by adjusting the input virus amount (10x) to ensure similar cell lysate luciferase levels (Figure 2C-D). With similar entry levels, we observed that the ∼300% increase in release efficiency was maintained and variability among trials was reduced (Figure 2C). In contrast, the release efficiency of Vesicular Stomatitis virus, another enveloped virus, was not affected by the lack of PS receptors on the surface of the cells (Figure 2E-F). We further confirmed the phenotype observed for CHIKV by omitting the viral entry step through the production of nano-luciferase tagged virus-like particles (VLPs) in the cells via plasmid transfection of a structural cassette. CHIKV VLPs release efficiency similarly displayed a 3-fold increase in VeroΔTIM/AXL cells (Figure 2G-H). Together these data indicate that cells lacking TIM release more CHIKV particles.

**Figure 2.**
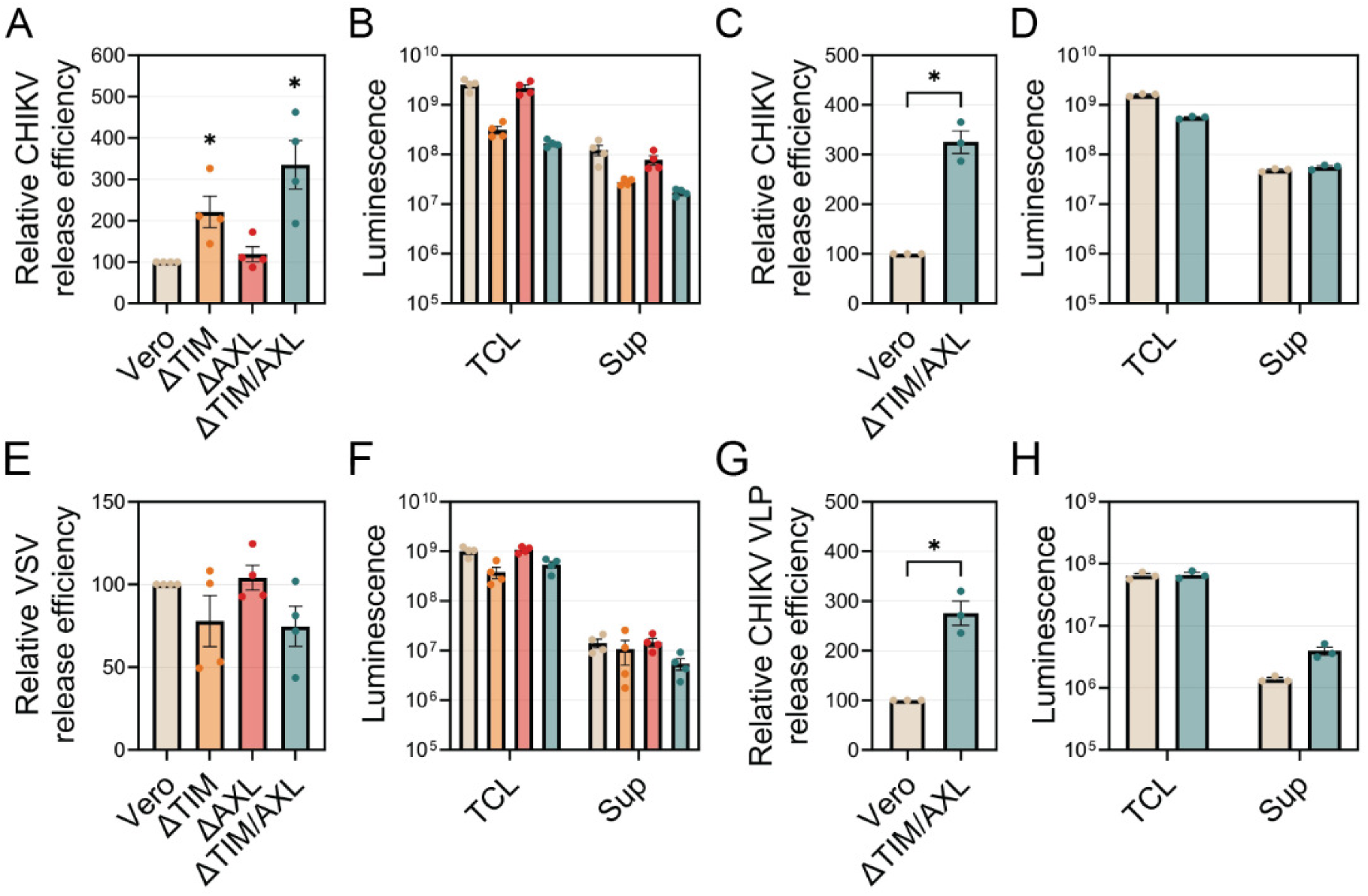
CHIKV particles are released more efficiently from Vero cells lacking TIM-1. Release efficiency assay of CHIKV-GFP-E2-NLuc in Vero, VeroΔTIM, VeroΔAXL, and VeroΔTIM/AXL cells (MOI 0.5, harvested at 18 hours) **(A)** and corresponding levels of luminescence present in the total cell lysates (TCL) and supernatants (sup) **(B)**. Release efficiency assay with VeroΔTIM/AXL cells infected with ten times more CHIKV-GFP-E2-NLuc than Vero cells **(C)** and corresponding luminescence levels **(D).** Release efficiency assay of VSV-GFP-M-NLuc in Vero, VeroΔTIM, VeroΔAXL, and VeroΔTIM/AXL cells (MOI of 1, harvested at 8hrs) **(E)** and corresponding luminescence levels **(F).** Release efficiency assay of CHIKV VLPs in Vero and VeroΔTIM/AXL cells transfected with a plasmid encoding CHIKV’s structural cassette tagged with nano-luciferase **(G)** and corresponding luminescence levels **(H).** Data represent the mean ±SEM from at least three independent trials. For each release assay, data was normalized to the parental cell line to determine the relative release efficiency. Unpaired parametric Student’s t-test with unequal variance (Welch’s correction) was performed to determine statistical significance in comparison to the parental cell line. *, p < .05.

### The presence of PS receptors increases levels of cell-associated virus through binding to envelope PS of budding virions

Our data in Vero cells suggested that TIM-1 limited CHIKV particle release to a greater extent than AXL. TIM-1 is an integral membrane protein with a structure comprised of an N-terminal globular domain, a long highly glycosylated stem region, a transmembrane domain, and a cytoplasmic tail. The globular N-terminal domain contains the PS binding site (18, 23, 24). We hypothesized that TIM-1 binding to CHIKV envelope-PS decreases virion release from infected cells. Therefore, we asked if we could promote particle release by saturating TIM-1 with PS liposomes post-virus entry. Fluorescently labeled PC:PE:PS liposomes were added to infected Vero or VeroΔTIM/AXL cells 6 hpi and release efficiency was calculated after 12 hours (Figure 3A). The addition of liposomes caused a dose-dependent increase in release efficiency in parental Vero cells but did not impact release in cells lacking PS receptors (Figure 3B). Exogenous expression of hTIM-1 in parental Vero cells did not result in significant changes in the release efficiency of CHIKV (Fig 3C). Vero cells naturally produce TIM-1 and transfection is unable to increase TIM-1 levels further (17). In contrast, transfection of exogenous hTIM-1 in VeroΔTIM/AXL cells significantly decreased the release efficiency of CHIKV (Figure 3D).

**Figure 3.**
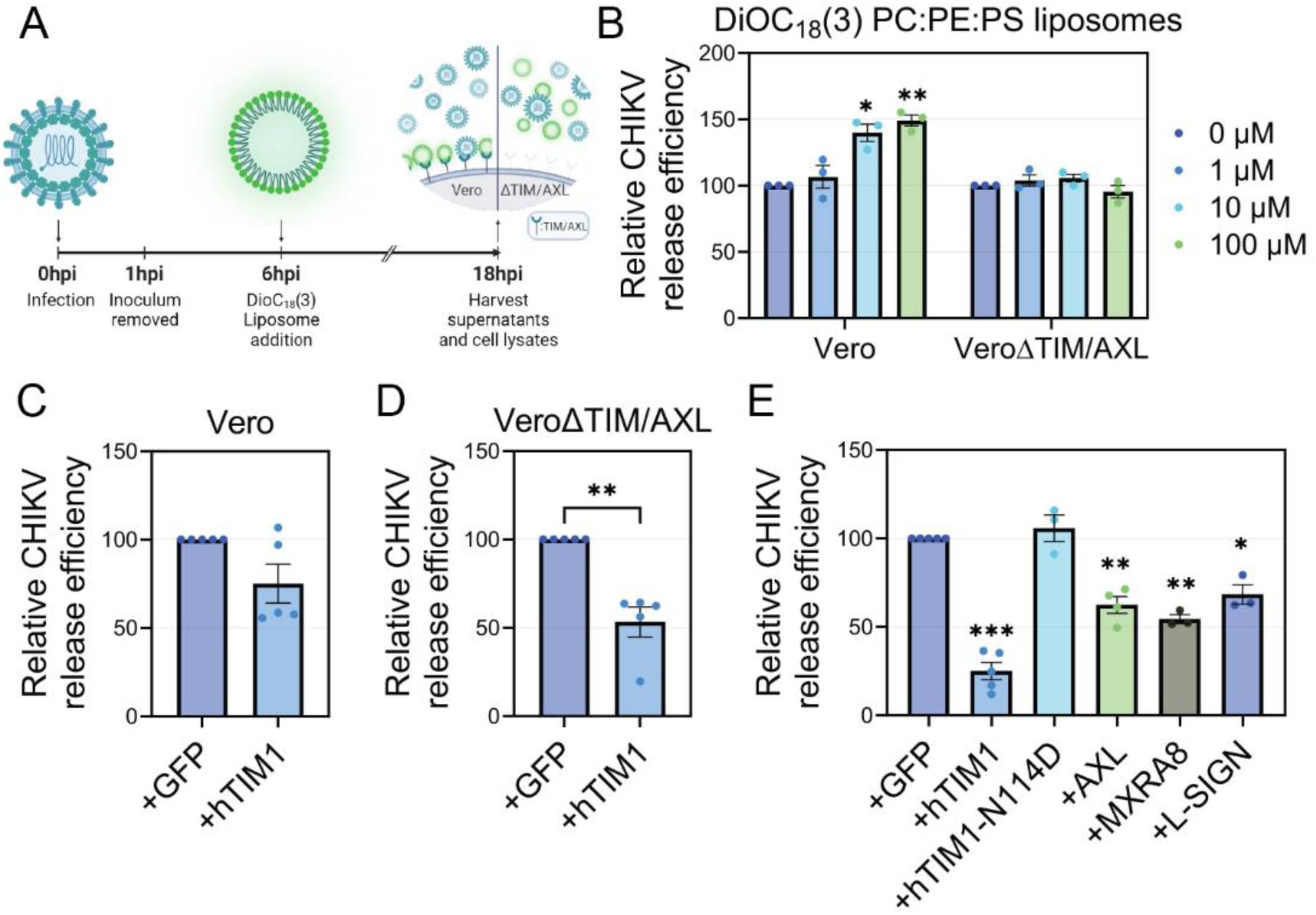
Chikungunya binds to the phospholipid binding domain of TIM-1, preventing its efficient release. **(A)** Experimental design for fluorescent liposome competition during release of CHIKV-infected Vero and VeroΔTIM/AXL cells. Created in BioRender.com **(B)** Increasing concentrations of fluorescent PC:PE:PS liposomes were added to CHIKV-E2-NLuc infected Vero or VeroΔTIM/AXL cells 6 hpi and release efficiency was calculated 18hrs post-infection. Data was normalized to the no-liposome control of each cell line. Vero **(C)** and VeroΔTIM/AXL **(D)** cells were transfected with a plasmid encoding hTIM-1 and infected with CHIKV-E2-NLuc 24 hours following transfection. Release efficiency was calculated 18hrs post-infection. **(E)** 293T cells were transfected with plasmids encoding the indicated surface receptors and infected with CHIKV-E2-NLuc 24 hours following transfection. Release efficiency was calculated 18hrs post-infection. Data was normalized to the GFP-transfected control. Data represents the mean ±SEM from at least three independent trials. Unpaired parametric Student’s t-test with unequal variance (Welch’s correction) was performed to determine statistical significance in comparison to the parental cell line. *, p < .05; **, p < .01; ***, p < .001.

Next, we examined viral release in 293T cells producing different molecules known to mediate CHIKV entry into mammalian cells (17, 19, 25, 26). 293T cells do not produce TIM-1, AXL, MXRA8 nor L-SIGN receptors (17, 27). Therefore, cells were transfected with plasmid expression vectors and release efficiency was compared to transfection of a plasmid encoding GFP for a control (Figure 3E). Similar to our previous data (17), production of TIM-1, MXRA8, and L-SIGN increased the entry efficiency of CHIKV as evidenced by the higher cell lysate luciferase activity (Supplemental Figure 1). TIM-1 production decreased particle release by ∼75%, while AXL, MXRA8, and L-SIGN decreased particle decrease by approximately 50%. Transfection of a TIM-1 mutant deficient in PS binding (N114D) (18) displayed similar release efficiency as GFP. These data suggest that CHIKV particle release can be suppressed by the overproduction of entry factors, although TIM-1 was the most efficient.

### Cellular knockout of CDC50a flippase subunit displays changes in Chikungunya virus entry, replication, and release efficiency

In our previous study, we produced PS-rich CHIKV particles by knocking out the flippase chaperone CDC50a, which increased outer leaflet PS in the plasma membrane of host cells (17). Unexpectedly, we observed phenotypic differences in CHIKV replication cycle in cells lacking CDC50a (ΔCDC50) that may also indicate enhanced particle release. In this study, we aimed to further investigate the relationship between outer leaflet PS and CHIKV using human haploid (HAP1) and vervet monkey kidney (VeroS) ΔCDC50 cells. CHIKV entered both HAP1 cell lines with similar efficiencies (Figure 4A). Yet, supernatant titers were consistently higher from CHIKV-infected HAP1ΔCDC50 cells than parental HAP1 cells (Figure 4B). CHIKV virions produced in HAP1ΔCDC50 cells contain higher levels of outer leaflet PS which results in higher particle specific infectivity when titrated on Vero cells (17). To determine if the higher titers observed in HAP1ΔCDC50 cells were all due to the enhanced particle infectivity, we examined the release efficiency from the cells. We observed a mild increase in CHIKV release in HAP1ΔCDC50 cells despite similar luminescence levels in the cell lysates (Figure 4C, Supplemental Figure 2A). This suggests that HAP1ΔCDC50 cells release more particles which are more infectious when compared to parental HAP1 cells. Interestingly, we observed decreased levels of surface TYRO3 in uninfected HAP1ΔCDC50 cells, which may enhance CHIKV release (Figure 4D). TYRO3 is the only known PS receptor produced in HAP1 cells. However, CHIKV does not rely solely on TYRO3 for entry in HAP1 cells (17), explaining why initial entry was not affected.

**Figure 4.**
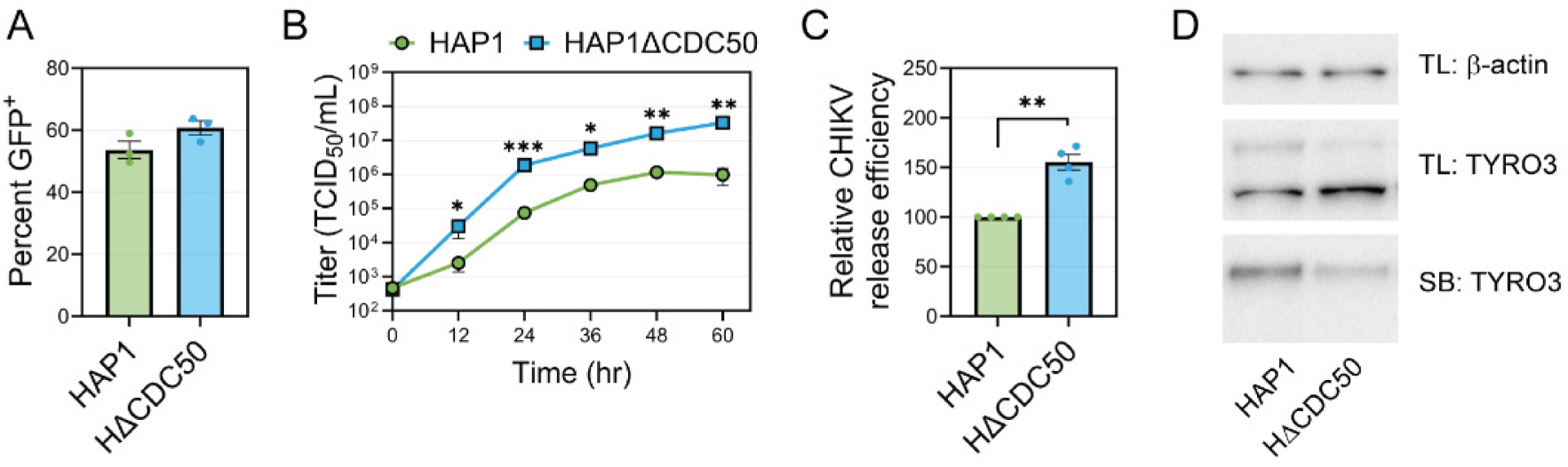
CHIKV displays an increase in release efficiency in HAP1ΔCDC50 cells and a decrease in surface receptor Tyro3. **(A)** Entry efficiency of CHIKV-GFP in HAP1 and HAP1ΔCDC50 cells. **(B)** Multi-cycle replication curve of CHIKV in HAP1 and HAP1ΔCDC50 cells (MOI 0.01). **(C)** Release efficiency of CHIKV-GFP-E2-NLuc in HAP1 and HAP1ΔCDC50. Data was normalized to the release efficiency of the parental cell line to determine the relative release efficiency. **(D)** Surface biotinylation analysis of uninfected HAP1 and HAP1ΔCDC50 cells. Total lysates and surface proteins were probed using a Tyro3 antibody or Actin antibody as a loading control. Data represents the mean ±SEM from at least three independent trials. Unpaired parametric Student’s t-test was performed to determine statistical significance in comparison to the parental cell line at each indicated timepoint. An unequal variance (Welch’s correction) t-test was performed for normalized data. *, p < .05; **, p < .01; ***, p < .001.

Unlike in HAP1ΔCDC50 cells, CHIKV entry was dramatically decreased in VeroSΔCDC50 cells (Figure 5A). While few CHIKV particles were able to initiate infection in the two-hour entry assay, if CHIKV infection was not limited, there was cell-to-cell spread detected (Figure 5B). CHIKV infection required an additional 24 hrs in VeroSΔCDC50 cells to obtain a similar number of GFP-positive cells (Figure 5B). Supernatant titers from VeroS cells were higher during the early time points in the multi-cycle replication curve, but by 48 hr CHIKV-infected VeroSΔCDC50 cells produced higher titers than parental VeroS cells (Figure 5C). A stronger increase in release efficiency was observed in VeroSΔCDC50 cells in comparison to the HAP1ΔCDC50 cells (Supplemental Fig 2B-C). To overcome the entry defect observed in VeroSΔCDC50 cells, we evaluated the release efficiency after infecting VeroSΔCDC50 with five times more virus than parental cells (Fig 5D). This led to similar levels of luminescence in the cell lysates of parental and VeroSΔCDC50 cells (Supplemental Fig 2D). VeroS cells lacking CDC50a activity produced 3-4 fold more CHIKV particles than parental VeroS cells (Figure 5D). Transfection of a plasmid encoding CDC50a in VeroSΔCDC50 cells significantly decreased the release efficiency of CHIKV (Figure 5E).

**Figure 5.**
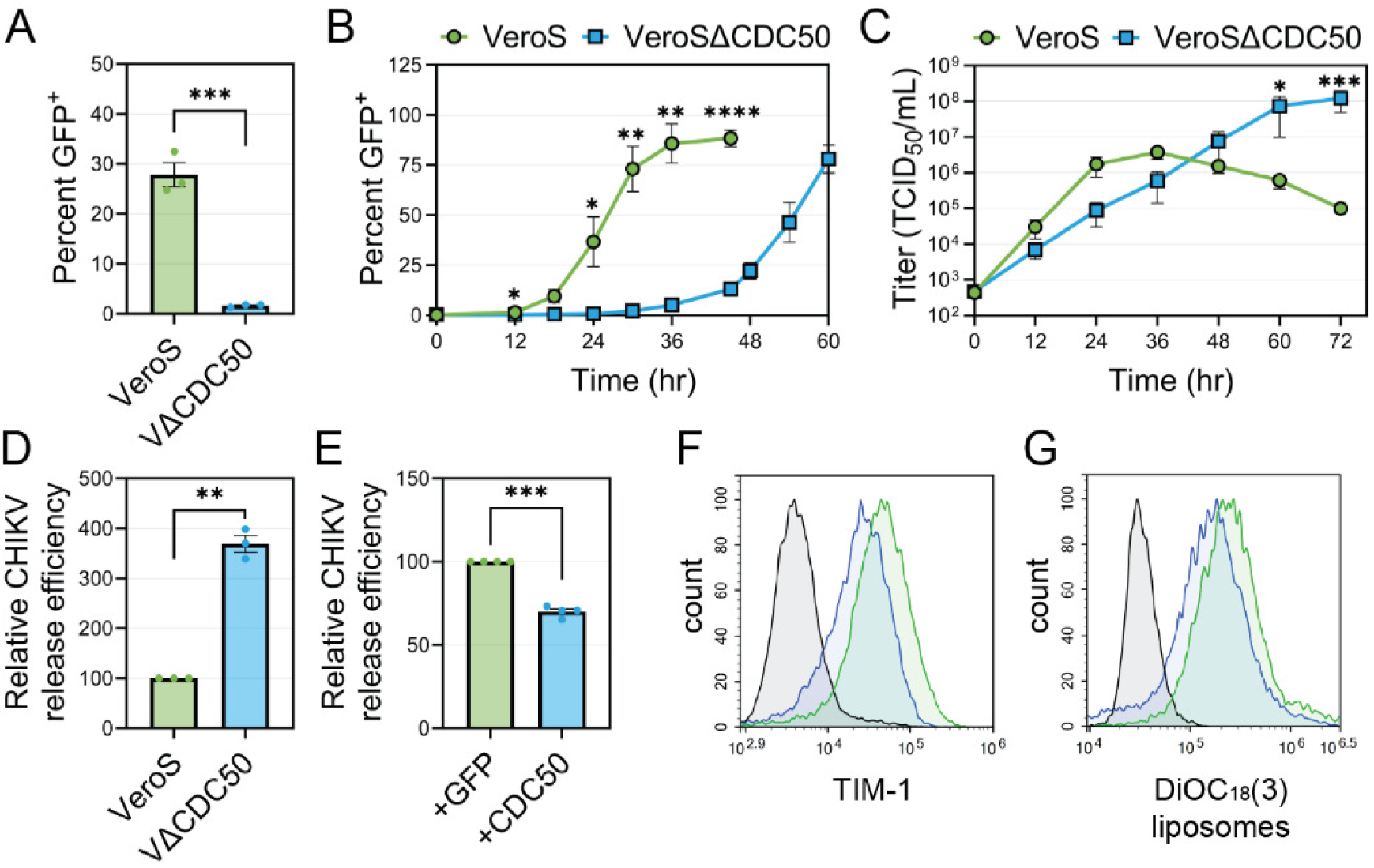
CHIKV entry is reduced, yet release is enhanced in VeroSΔCDC50 cells. **(A)** Entry efficiency of CHIKV-GFP in VeroS and VeroSΔCDC50 cells. **(B)** CHIKV-GFP spread in VeroS and VeroSΔCDC50 cells (MOI 0.1). **(C)** Multi-cycle replication curve CHIKV in VeroS and VeroSΔCDC50 cells (MOI 0.01). Supernatants were harvested and titrated at the indicated time points. **(D)** Release efficiency assay of CHIKV-GFP-E2-NLuc when five times more virus as added to VeroSΔCDC50 than Vero cells to equalize cell lysate luminescence levels. **(E)** VeroSΔCDC50 cells were transfected with a plasmid encoding CDC50a and infected with CHIKV-E2-NLuc, release efficiency was assessed 18 hours post-infection. Surface receptors of VeroS (green) and VeroSΔCDC50a (blue) were assessed via staining using **(F)** a TIM-1 antibody or **(G)** binding of DioC_18_(3) fluorescent PC:PE:PS liposomes and analyzed through flow cytometry. Data represents the mean ±SEM from at least three independent trials. Unpaired parametric Student’s t-test was performed to determine statistical significance in comparison to the parental cell line at each indicated timepoint. An unequal variance (Welch’s correction) t-test was performed for normalized data. *, p < .05; **, p < .01; ***, p < .001; ****, p < .0001.

CHIKV entry into VeroS cells is dependent on TIM-1 (17). Because VeroSΔCDC50 cells display altered PS distribution, we hypothesized that the surface levels of TIM-1, a PS receptor, might be affected. When we examined surface TIM-1 production in uninfected VeroSΔCDC50 cells by surface staining (Figure 5F) and binding of fluorescently labeled PC:PE:PS liposomes (Figure 5G), we noted a decrease that may explain both the decrease in CHIKV entry into VeroSΔCDC50 cells and the enhanced CHIKV release phenotype.

### CHIKV release efficiency correlates with the presence of phospholipid-binding receptors across cell lines

To examine the correlation between cell surface PS receptors and release efficiency, we compared a panel of cell lines. To evaluate the presence of phospholipid binding receptors across cell lines without species-specific antibodies, we used fluorescently labeled PC:PE:PS liposomes and quantified cellular binding through flow cytometry (Figure 6A-B). We then assessed CHIKV particle release in each cell line (Figure 6C, Supplemental Figure 3). Surprisingly, CHIKV displayed the highest levels of release in BSR-T7/5 cells. Similar findings were previously evidenced with BHK cells in Ramjag, et al., 2022 (22). We observed an inverse correlation between PC:PE:PS liposome binding and particle release in Vero, VeroS, VeroΔTIM/AXL, Aag2, and BSR-T7/5 cells (Figure 6D). The release efficiency in mosquito cell lines, C6/36 and Aag2, was similar to that of Vero cells despite not having any identified TIM or AXL homologs. Surprisingly, Aag2 cells displayed liposome binding levels two-fold above background strongly suggesting the presence of PC:PE:PS binding receptors in these cells. This data further demonstrates the strong effect that phospholipid binding receptors have on the release of CHIKV virions.

**Figure 6.**
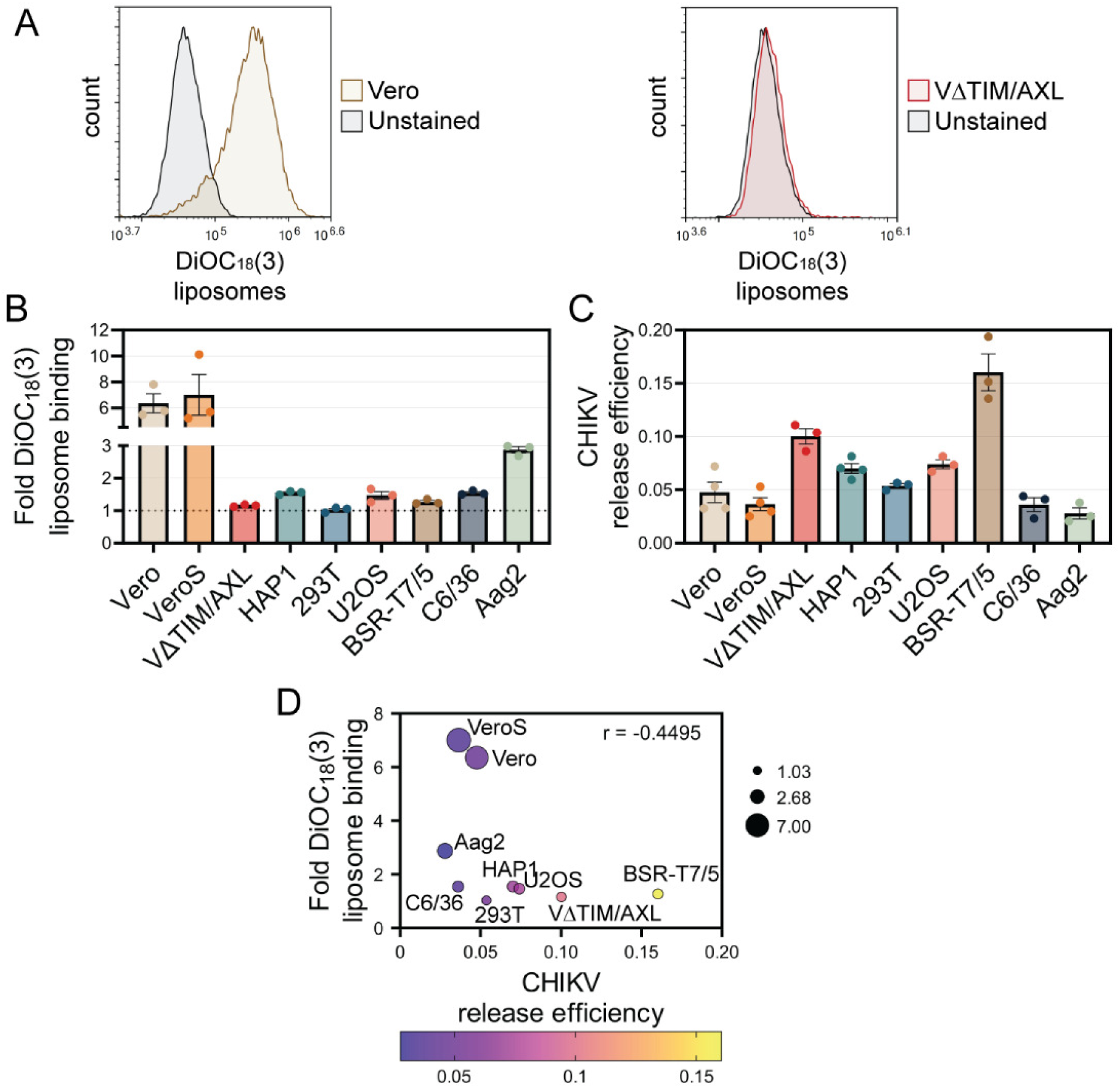
CHIKV release efficiency correlates with PC:PE:PS liposome binding in a panel of cell lines. **(A)** Representative histograms of binding of fluorescent liposomes in Vero and VeroΔTIM/AXL cells as a measure for phospholipid binding receptors. **(B)** Fold binding of fluorescent PC:PE:PS liposomes in mammalian and mosquito cell lines. To remove differences in fluorescent background levels among cell lines, fold binding was determined by calculating the ratio of DioC_18_(3) mean fluorescent intensity (MFI) over no-liposome control for each cell line. The dotted line represents the threshold where DioC_18_(3) MFI was equivalent to no-liposome background levels indicating no binding occurred. **(C)** The release efficiency of CHIKV-GFP-E2-NLuc in a panel of mammalian and mosquito cell lines. **(D)** Correlation analysis between liposome binding and CHIKV release efficiency. The size of circles represents the degree of liposome binding and colors indicate levels of release efficiency. Data represents the mean ±SEM from at least three independent trials.

### TIM-1 is downregulated from the cell surface following CHIKV infection

To enhance particle release and prevent superinfection, many viruses downregulate viral receptors (28–30). This downregulation can be through receptor saturation and subsequent endocytosis or direct receptor degradation (21, 31). For example, Japanese encephalitis virus (JEV) counteracts AXL’s viral release inhibition by inducing AXL degradation through the ubiquitination pathway (21). Alternatively, HIV encodes an accessory protein, Nef, which induces the engulfment of TIM, reducing TIM protein from the cell surface (31). To examine if TIM-1 levels are changed by CHIKV infection, we infected VeroS cells and monitored TIM-1 levels on the plasma membrane. CHIKV infection decreased TIM-1 levels (Figure 7A) and the ability of cells to bind PC:PE:PS liposomes (Figure 7B). To determine if this difference was specific to TIM-1, cells were mock infected, infected with CHIKV-GFP or with Lymphocytic Choriomeningitis virus (LCMV)-GFP, and surface proteins were labeled with biotin when 90% of the cells were positive for GFP production (Figure 7C). Total cell lysates and purified surface biotinylated proteins were separated on an SDS-PAGE gel and visualized on a BioRad stain-free gel (Figure 7D). We found few differences between mock and infected cells, except for the production of the CHIKV envelope protein which was enriched in the surface fraction (Figure 7D, Supplemental Figure 4). This suggests that CHIKV infection does not cause a global decrease in the production of surface proteins. Yet, immunoblot analysis displayed an 85% reduction of TIM-1 levels in purified surface proteins, while transferrin and AXL levels decreased by ∼40% in CHIKV-infected cells (Figure 7E). Infection with LCMV only decreased TIM-1 and AXL surface levels by ∼15% and ∼10%, respectively, and increased levels of transferrin by ∼40% (Figure 7E).

**Figure 7.**
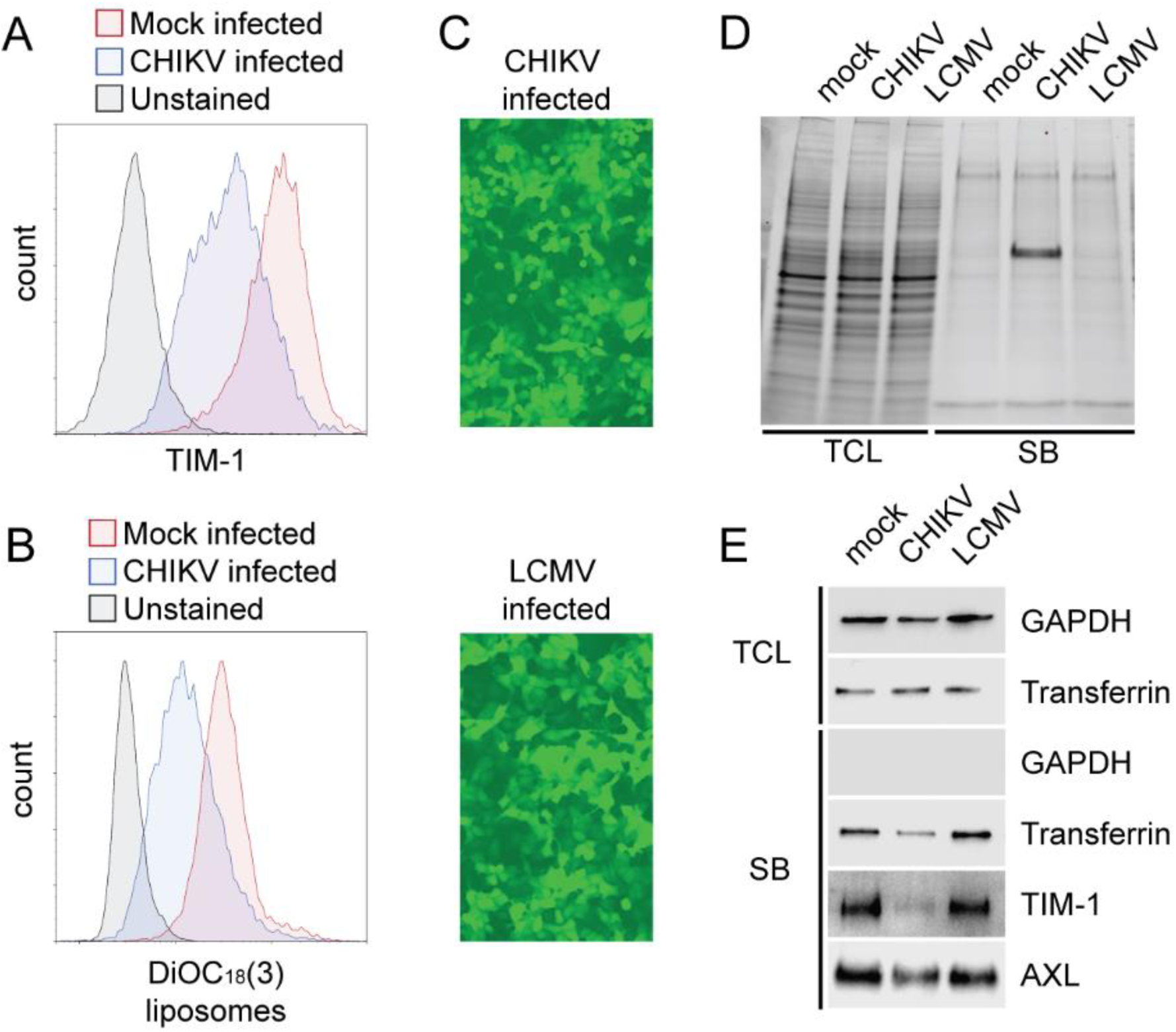
CHIKV infection decreases surface levels of TIM-1. Levels of TIM-1 in the surface of uninfected or CHIKV-infected VeroS cells were assessed via receptor staining using a TIM-1 antibody **(A)** or binding of fluorescently labeled liposomes **(B)** and analyzed through flow cytometry. **(C)** VeroS cells were infected with either CHIKV-GFP (top, MOI 0.5) or LCMV-GFP (bottom, MOI 1) resulting in similar levels of infection. **(D)** The total protein present in total cell lysates (TCL) and biotinylated surface proteins (SB) of uninfected, CHIKV, and LCMV-infected VeroS cells were compared using a stain-free gel. **(E)** Immunoblot analysis of samples shown in panel D.

To further understand the mechanism of surface TIM-1 downregulation we evaluated the timing of this phenotype. Cell surface TIM-1 levels were first reduced around 6-9hpi, concurring with E1 detection (Figure 8A Supplemental Figure 5A-B), and continued to decrease over time. After 12-15hr after infection, we observed a decrease of ∼50% in the binding of fluorescently labeled liposomes (Figure 8B-C). To determine if a specific CHIKV protein triggers the decrease in surface TIM-1 levels, we transfected VeroS cells with plasmids encoding each of CHIKV’s proteins. Production of each viral protein was confirmed through immunoblots (Supplemental Figure 5C-D). We observed a decrease in PC:PE:PS liposome binding after the production of CHIKV nsP2 (Figure 8D). This data suggests that the decrease of TIM-1 in infected cells might be mediated through the viral protease nsP2.

**Figure 8.**
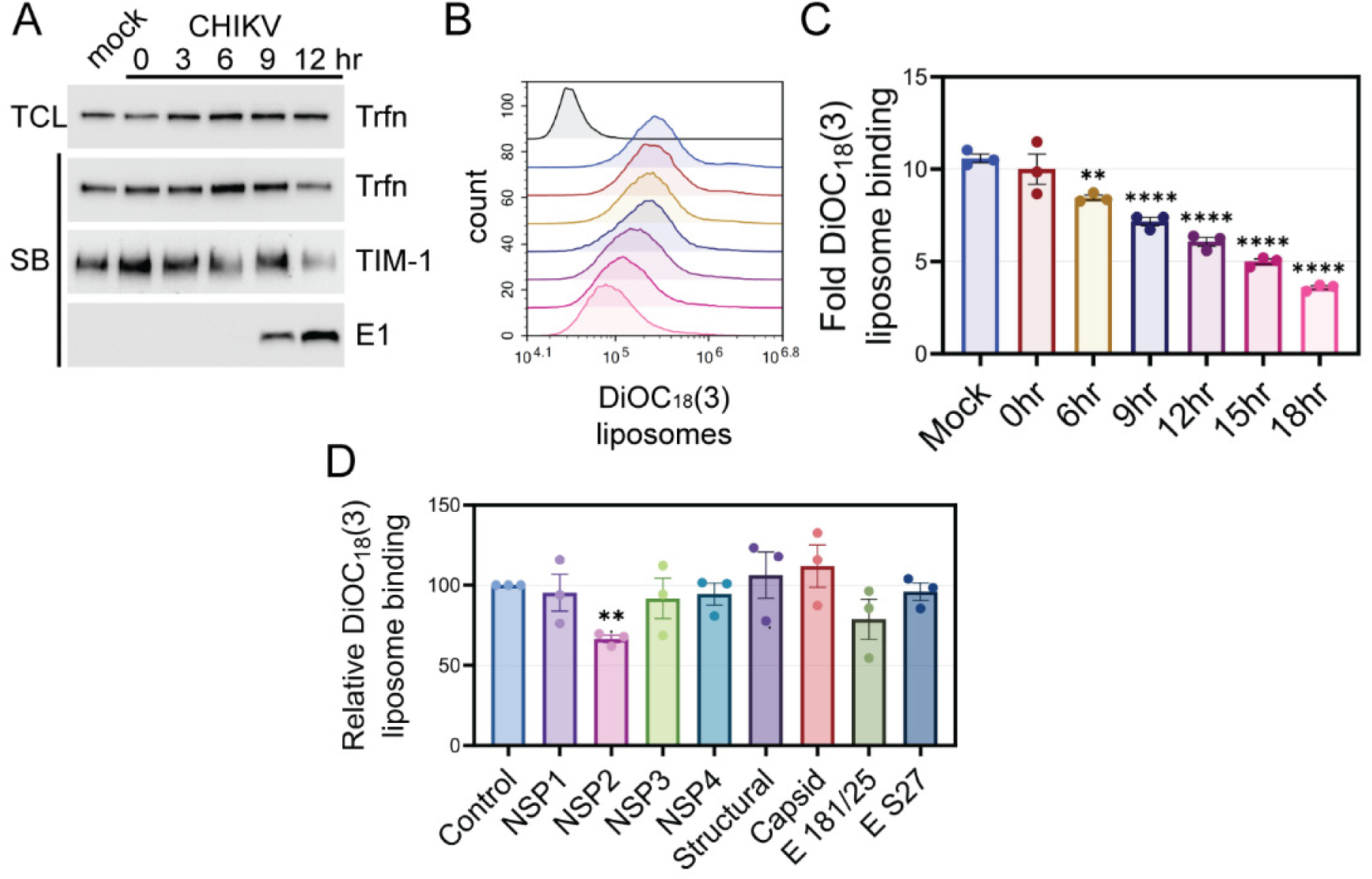
Levels of surface TIM-1 start decreasing around 9hpi in a mechanism possibly mediated by nsP2. **(A)** CHIKV-GFP infected VeroS cells were infected at different time points, and all were harvested at the same time for surface biotinylation analysis. Infection was maintained for 0, 3, 6, 9 or 12 hours. Samples were probed using TIM-1, E1, or transferrin (Trfn) antibodies. **(B)** VeroS cells were infected with CHIKV at different time points, and subjected to fluorescent liposome binding simultaneously. Infection was maintained for 0, 3, 6, 9, 12, 15, or 18 hours and analyzed through flow cytometry. **(C)** Quantification of fluorescent PC:PE:PS liposomes. **(D)** Cells were transfected with plasmids encoding CHIKV proteins for 48 hours and analyzed through binding of fluorescently labeled liposomes using flow cytometry. Liposome binding was compared to control cells to determine the relative liposome binding levels. Data represents the mean ±SEM from at least three independent trials. An ordinary one-way ANOVA with multiple comparisons was used to evaluate statistical differences in comparison to control. An unequal variance (Welch’s correction) t-test was performed for normalized data. *, p < .05; **, p < .01; ****, p < .0001.

## DISCUSSION

Our study provides evidence that surface receptors can prevent efficient CHIKV viral release. TIM-1 appeared to be more effective than TAM family receptors (*i.e.,* AXL) and other entry factors (*i.e.,* MXRA8 or L-SIGN) at preventing virions from completing their egress from infected cells. We propose that the release inhibition observed in Vero cells is mediated through the interaction between the PS binding domain of TIM-1 and the lipid envelope surrounding CHIKV particles (Figure 9). CHIKV entry is efficiently mediated by different molecules depending on the cell line (17). Presumably, these same factors that mediate entry can also capture newly formed particles, ultimately reducing release. When various entry factors were transfected into 293T cells we observed they each reduced release, and TIM-1 was most effective. While most of the work presented here focused on Vero cells and TIM-1, the main entry receptor for CHIKV in these cells, we hypothesize removal of entry receptors important to other cell types would also enhance CHIKV release.

**Figure 9.**
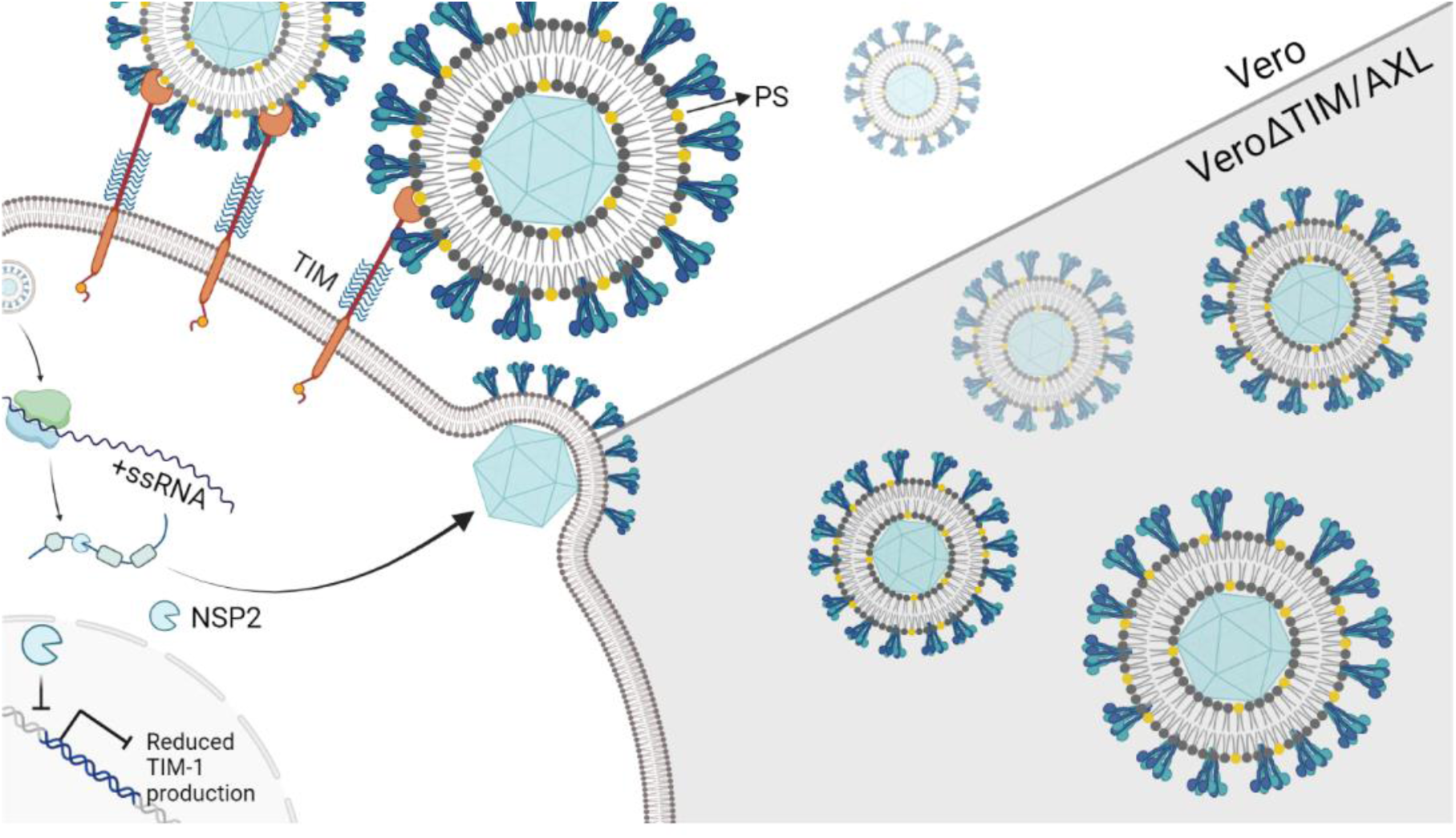
CHIKV release is decreased by TIM-1 binding to envelope PS. **(A)** Diagram of mechanistic model displaying budding virions attached to TIM-1 in Vero cells and being released efficiently from VeroΔTIM/AXL cells. Nonstructural protein 2 (nsp2) decreases levels of TIM-1 over time possibly through cellular transcription shutoff. Created in BioRender.com

PS receptors from the TIM and TAM families interact with PS differently, which may contribute to phenotypic differences observed in viral release. Receptors from the TAM family, including TYRO3 and AXL, require a bridging ligand known as Gas6 (32). Previous studies demonstrate this cofactor is present in the fetal bovine serum supplemented in the media of tissue culture cells at concentrations typically required to bridge cell-PS binding (33). Although our infections took place in serum-containing media, AXL and/or Gas6 levels may not have been sufficient to link newly formed particles to the cell surface as well as TIM-1. CHIKV particles are made up of a highly organized lattice of glycoproteins with limited access to the lipid layer (2). Gas6 may not be able to access CHIKV PS as well as TIM-1 limiting the TAM family’s ability to both mediate entry and reduce particle release. While we did not find a role for AXL in limiting CHIKV release in Vero cells, it is important to recognize that AXL can inhibit the release of other viral particles, as is the case for Japanese Encephalitis virus (JEV) (21).

HIV, JEV, and Ebola virus release is limited by PS receptors (20, 21). We found that particle retention by TIM and AXL could significantly reduce CHIKV release, but was not able to significantly reduce Vesicular Stomatitis virus (VSV) release. VSV infection consistently produces higher titers after a single round of infection compared to CHIKV, this may enable VSV to quickly saturate the PS binding sites and produce enough particles that limit the ability to observe a release defect. This phenotype could be general for a wider variety of enveloped viruses and may suggest viruses that produce fewer virions per cell may be impacted more than viruses that produce larger quantities of particles.

We observed that increased release efficiency correlated with decreases in levels of surface receptors not only in VeroΔTIM/AXL but also in cells knocked out for the flippase subunit CDC50a. Cells knocked out for CDC50a lack flippase activity resulting in increased levels of outer leaflet PS, possibly leading to failure in efficient redistribution of their lipids to accommodate integral proteins. In general, transmembrane proteins (*e.g.,* TIM, AXL, TYRO3) can disrupt fluidity within the plasma membrane which can trigger changes in the translocation of specific lipids (34). Additionally, the composition of the plasma membrane can prevent the insertion of receptors into the bilayer and induce changes in their topological orientation (34–36). TIM proteins have been shown to preferentially enter the lipid bilayer among unsaturated phospholipids rather than saturated ones (35). Consequently, we hypothesize that ΔCDC50 cells might undergo a redistribution of membrane proteins and a decrease in the proper insertion of these membrane receptors (*i.e.,* TIM and TYRO3), resulting in increased CHIKV release.

CHIKV infection decreased cell surface TIM-1 in a mechanism possibly mediated by nsP2 protease. While infection also reduced surface proteins AXL and transferrin, surface TIM levels were more significantly depleted. nsP2 shuts off cellular transcription and has been suggested to be one of the main factors behind superinfection exclusion in alphaviruses by interfering with the formation of replication complexes of incoming viruses (37–39). Therefore, the CHIKV-induced receptor decrease may not be specific, but might disproportionally affect surface proteins with shorter half-lives such as TIM (half-life <2hrs) (31).

CHIKV release efficiency among cell lines correlated with the presence of phospholipid binding receptors. We found that the release efficiency of mosquito cells C6/36 and Aag2 was similar to that of Vero cells, which express TIM-1 and AXL receptors. Aag2 cells were able to significantly bind PS-containing liposomes, although previous studies have failed to identify homologs for PS receptors in mosquito cells. Future studies should further explore cellular receptors that might be playing a role in preventing the efficient exit of viral particles in these cells. Mosquito cells display potential budding of alphaviruses from internal membranes such as cytopathic vacuoles (40). It would be interesting if PS receptors or other cellular proteins present in these vacuoles could attach to new virions before they reach the cell surface. This mechanism would not be surprising as viruses such as JEV that bud from the endoplasmic reticulum have been shown to bind to AXL (21).

This study provides evidence for the importance of PS receptors during the egress of CHIKV. The ability of CHIKV to counteract this inhibition might result in higher levels of disease spread inside the host’s body. However, the significant increase in the production of viral particles from cells lacking TIM-1 and increased infectivity of virions previously observed in ΔCDC50 cells (17) could be employed to maximize particle production during vaccine development. Further studies should characterize the extent to which PS receptors could inhibit the efficient egress of other highly pathogenic enveloped viruses.

## MATERIALS AND METHODS

### Cells

All mammalian cell lines were maintained at 37°C and 5% CO_2_. Parental monkey Vero cells and Vero cells knocked out for TIM (VeroΔTIM), AXL (VeroΔAXL), and both (VeroΔTIM/AXL) were a gift from Dr. Wendy Maury from the University of Iowa (41). All Vero cells, including Vero-humanSLAM (VeroS) (42) and VeroS knocked out for CDC50a chaperone (VeroSΔCDC50) (17), and BHK stably expressing T7 RNA polymerase (BSR-T7/5) (43) cells were maintained in DMEM supplemented with 5% FBS. Parental human haploid cells (HAP1) and HAP1 knocked out for CDC50a (HAP1ΔCDC50) cells were purchased from Horizon Discovery and maintained in Iscoves’ modified Dulbecco’s Medium (DMEM) supplemented with 8% fetal bovine serum (FBS). Human 293T cells and human osteosarcoma U2OS, a gift from Dr. Neale Ridgway from Dalhousie University (44), were maintained with DMEM media supplemented with 10% FBS. Mosquito cell lines were kept at 28°C and maintained in Leibovitz’s L-15 media supplemented with 10% FBS (C6/36 – *Aedes albopictus*) or HyClone SFX-Insect media supplemented with 2% FBS (Aag2 - *Aedes aegypti*).

### Viruses

All Chikungunya infections were performed using the attenuated vaccine strain 181 clone 25 (181/c25). Full-length CHIKV genome was untagged (CHIKV), encoded *gfp* as an additional transcription unit between the non-structural and structural gene (CHIKV-GFP) (45) or contained NLuc inserted at the N-terminus of E2 (CHIKV-E2-NLuc, CHIKV-GFP-E2-NLuc). The described changes were introduced into the molecular clone pSinRep5-181/25c (Addgene cat. 60078), a gift from Dr. Terrance Dermody. To recover the virus, plasmids were linearized and *in vitro* transcribed with the mMessage mMachine SP6 transcription kit (Invitrogen, cat. AM1340) per the manufacturer’s protocol to produce the full-length positive-sense mRNA which was transfected into Vero cells using Lipofectamine 3000 following the manufacturer’s instructions. Vesicular Stomatitis virus (VSV) used to perform release efficiency assays was tagged with nano-luciferase in the coding region of the matrix protein (M) following residue 37 and encodes GFP as an additional transcriptional unit at a post-G site (VSV -M-NLuc-GFP) as described in (46, 47). Tri-segmented attenuated lymphocytic choriomeningitis virus encoding GFP (LCMV-GFP) was a gift from Dr. Luis Martínez-Sobrido (48). CHIKV and VSV stocks were propagated using Vero cells and LCMV stocks were propagated in BSR-T7/5 cells. All stocks were titrated on Vero cells using serial dilutions to determine the tissue culture infection dose 50 (TCID50) according to the Spearman-Karber method.

### Virus Release Assays: Immunoblots

Vero or VeroΔTIM/AXL cells were plated in 10 cm^2^ dishes at a density of 2.5×10^6^ per plate, one day before infection. Cells were infected for one hour with CHIKV-GFP at an MOI of 0.5 (Vero, ΔTIM/AXL 1x) or 5 (ΔTIM/AXL 10x) and incubated at 37°C. Eighteen hours following infection, supernatants were collected, and cells were lysed in M2 lysis buffer (50 mM Tris [pH 7.4], 150 mM NaCl, 1 mM EDTA, 1% Triton X-100) for 5 minutes and collected. Cell lysate samples were cleared by centrifuging at 6,000xg for 25 minutes. Supernatant samples were cleared twice at 6,000xg for five minutes and concentrated with ultracentrifugation over a 20% sucrose cushion for 3 hours at 28,000 rpm at 4°C. Purified pellets were resuspended in 200μl of 1x PBS. Cell lysates and purified supernatants were separated on an SDS-PAGE and analyzed through immunoblotting against vinculin as a loading control (1:2,000, MGA465GA, BioRad) or CHIKV E1 glycoprotein (1:1,000, MAB97792, R&D systems). Protein levels were quantified through Image Lab 6.1 densitometry analysis.

### Multi-step Replication Curves

Vero, VeroS, and VeroSΔCDC50 cells were plated at 2.5×10^5^ cells/well in a 12-well plate while HAP1 and HAP1ΔCDC50 cells were plated at a density of 3.0×10^5^ cells/well. Cells were infected with untagged CHIKV or CHIKV-GFP-E2-NLuc at an MOI of 0.01 for one hour. At each indicated time point, supernatants were collected and replaced with corresponding media containing FBS. Samples were titrated by calculating the tissue culture infection dose 50 (TCID50) on Vero cells using the Spearman-Karber method. Luminescence was quantified using the Nano-Glo Substrate (Promega) and measured in a GloMax Explorer (Promega) luminometer.

### Virus Release Assays: Luminescence

Cells were plated in 24-well plates at a density of 2.5×10^5^ cells/well, one day before infection. CHIKV inoculum was added for one hour at an MOI of 0.5 unless stated otherwise and incubated at 37°C. 18 hours following infection, supernatants were collected, and cells were lysed in M2 lysis buffer for 5 minutes and collected. Samples were cleared by centrifuging at 17,000xg for either 5 minutes (supernatants) or 25 minutes (cell lysates). Luminescence in supernatants and cell lysates was determined using the Nano-Glo Substrate (Promega) and measured in a GloMax Explorer (Promega) luminometer. Viral release was calculated as the ratio of luminescence in the supernatant divided by the luminescence in the cell lysates. Vesicular Stomatitis virus infections were performed at an MOI of 1 for one hour and samples were harvested 8 hours post-infection as described above.

### Virus Release Assays: Transfections and Plasmids

Vero and VeroΔTIM/AXL cells were plated in a 24-well plate at a density of 5×10^4^ cells/well. The following day, cells were transfected with a plasmid encoding CHIKV’s structural cassette (C, E3, E2, 6K, E1) with NLuc inserted at the N-terminus of E2 to produce luminescence viral-like particles (VLPs). Transfections were performed using Jet Optimus (Polyplus, #101000025) following the manufacturer’s protocol. Supernatants and cell lysates were collected 24 hours post-transfection and release assays were performed as described above.

Vero and VeroΔTIM/AXL cells were also transfected with plasmids encoding hTIM-1-GFP or GFP using Jet Optimus. 24 hours following transfection, CHIKV inoculum was added at an MOI of 0.5 for one hour and supernatants and cell lysates were harvested 18 hours post-infection. Release assay was performed as described above.

293T cells were plated in a 24-well plate at a density of 1.5×10^5^ cells/well one day before transfections. The following day, cells were transfected with plasmids encoding hTIM-1-GFP, TIM-1-N114D, AXL, MXRA8, or L-SIGN using Jet Prime (Polyplus, #101000027) following the manufacturer’s protocol. TIM plasmids were a gift from Dr. Wendy Maury (18). AXL (BC032229), MXRA8 (BC006213) (17), and L-SIGN (BC038851) plasmids were purchased from Transomic and if necessary, cloned into expression vectors. The following day, cells were infected, and release assays were performed as described above.

### Virus Release Assays: DioC_18_(3) PC:PE:PS liposomes

PC:PE:PS liposomes (75% PC: 20% PE: 5% PS) were prepared as described in (49) with the addition of DiOC_18_(3) (3,3’-Dioctadecyloxacarbocyanine Perchlorate) (Invitrogen, D275) for fluorescence, following manufacturer’s indications. Vero and VeroΔTIM/AXL cells were plated at a density of 2.5×10^5^ cells/well in a 24-well plate. The next day, cells were infected at an MOI of 0.5 for one hour. Six hours post-infection, DioC_18_(3) PC:PE:PS liposomes were added to the cells at the indicated concentrations. Supernatants and cell lysates were collected 18 hours post-infection and release assays were performed as described above.

### Entry Assays

Cells were plated in a 48-well plate at a density of 1×10^5^ cells/well (HAP1, HAP1ΔCDC50) or in a 24-well plate at 1.25×10^5^ cells/well (VeroS, VeroSΔCDC50). The next day, cells were infected with enough CHIKV-GFP infectious viral particles to obtain approximately 50% of infected cells after 12 hours. Inoculum was removed from the cells after one hour and treated with 30 mM ammonium chloride (NH_4_Cl) after 2 hours to prevent subsequent rounds of infection. Infected cells were resuspended in PBS, fixed using 4% formaldehyde, and the percentage of GFP+ cells was quantified using a NovoCyte Quanteon cytometer (Agilent).

### Surface Biotinylation

HAP1 and HAP1ΔCDC50 cells were plated in a 6-well plate at a density of 1×10^6^ cells/well. Cells were either mock infected or infected one day after plating with CHIKV-GFP (MOI 0.5) or LCMV-GFP (MOI 1) and harvested at the indicated time points. Cells were washed with cold PBS, and surface proteins were biotinylated with 0.5 mg/mL sulfosuccinimidyl-2-(biotinamido) ethyl-1,3-dithiopropionate (ThermoFisher) on ice for 45 minutes with gentle shaking. The reaction was quenched using Tris-HCl and cells were lysed with M2 lysis buffer at 4°C. Samples were centrifuged at 17,000xg for 10 minutes. A fraction of the lysate was saved (total cell lysate, TCL), and the surface proteins were bound to streptavidin Sepharose beads overnight at 4°C (GE Healthcare). Beads were then washed with buffer containing 100 mM Tris, 500 mM lithium chloride, 0.1% Triton X-100 followed by a buffer containing 20 mM HEPES [pH 7.2], 2 mM EGTA, 10 mM magnesium chloride, 0.1% Triton X-100. Samples were then analyzed through immunoblotting probing against TYRO3 (1:1000, R&D Systems, MAB859100), TIM (TIM (1:500, AF1750, R&D Systems), AXL (1:100, AF154, R&D Systems), GAPDH (1:2000, Santa Cruz Biotech, #sc-47724), Transferrin (1:1,000, PA5-27739, ThermoFisher), or CHIKV E1 (1:1,000, MAB97792, R&D systems).

### Cell-to-cell Spread Kinetics

VeroS and VeroSΔCDC50 were plated in a 24-well plate at a density of 1.25×10^5^ cells/well. One day after plating, cells were infected with CHIKV-GFP at an MOI of 0.1 for one hour. At the indicated time points, cells were lifted using trypsin, resuspended in PBS, and fixed in 4% formaldehyde. A NovoCyte Quanteon cytometer (Agilent) was used to analyze 10,000 live cells and quantify the percentage of GFP+ cells over time.

### Surface Receptor Staining

Cells were plated at a density of 1.0×10^6^ cells/well in a 6-well plate one day prior to staining. Cells were harvested either uninfected or after infection with CHIKV-GFP at an MOI of 0.5, 18 hours post-infection. Monolayers were cooled, washed, and treated with a blocking solution (dPBS +Ca2 +Mg2 with 2% (v/v) FBS) containing an anti-hTIM1(1:50-1:100, AF1750, R&D Systems) antibody. Samples were incubated at 4°C with gentle shaking for one hour and washed three times with ice-cold PBS. A blocking solution containing the corresponding secondary antibody, donkey anti-goat Alexa Fluor 594 (1:2500, A32758, Invitrogen), was added. Samples were incubated at 4°C in the dark with gentle shaking for 30 minutes. Samples were washed three additional times with PBS, lifted via scraping, and analyzed using a NovoCyte Quanteon cytometer (Agilent). Populations of live, single cells were gated using FSC/SSC and SSC-A/SSC-H, respectively. The GFP gate was set using uninfected, GFP-cells, and the AF594 gate was set with a secondary-only control. The AF594 MFI of 10,000 live, single, GFP+ cells was quantified. A 488-nm laser with a 530/30 “FITC” bandpass filter was used to assess GFP fluorescence, and AF594 was measured with a 561-nm laser with a 615/20 “PE-Texas Red” bandpass filter; all filter sets had default gain.

### Liposome binding assay

For comparison of liposome binding among different cell lines, cells were plated in a 12-well plate at a density of 5×10^5^ cells/well, and binding was assessed the following day. For assessing liposome binding following infection, VeroS cells were plated at a density of 2.5×10^5^ cells/well in a 12-well plate. CHIKV inoculum was added at an MOI of 0.5 for 1hr and binding was assessed after 18hrs. To evaluate the effect of CHIKV’s proteins on liposome binding, VeroS cells were plated in a 24-well plate at a density of 1×10^5^ cells/well. The following day, cells were transfected with plasmids encoding for CHIKV’s non-structural proteins with a FLAG tag (nsP1, nsP2, nsP3, and nsP4), a structural cassette (C, E3, E2, 6K, E1), capsid, E 181/25 (Southeast Asian strain) or E S27 (African strain). Plasmid encoding E S27 was a gift from Dr. Graham Simmons (50). Transfections were performed using Jet Optimus (Polyplus, #101000025) following the manufacturer’s protocol. Two days following transfection, binding was assessed.

To measure liposome binding, cells were placed on ice for 30 minutes. DioC_18_(3) PC:PE:PS liposomes were sonicated for 1 hour and added to the cells at a final concentration of 10µM. Liposomes were bound to cells for 1 hour on ice, removed, and washed with FBS-free media. Cells were lifted in FBS-free media and fixed in equal volume of 4% formaldehyde.

Samples were analyzed using a NovoCyte Quanteon cytometer (Agilent). Populations were gated using SSC-H/FSC-H and SSC-A/SSC-H to identify live and single cells, respectively. A 488-nm laser with a 530/30 “FITC” bandpass filter was used to assess DiOC_18_(3) fluorescence. A DiOC_18_(3)+ gate was set using non-liposome-treated cells as a DiOC_18_(3)-control. The DiOC_18_(3) MFI of 10,000 live, single events was quantified.

### Statistical Analysis

All graphs were made and analyzed using GraphPad Prism (v10.1.1, macOS). An unpaired parametric student’s T-test was performed to determine the significance between two groups. For data determining statistical significance among two groups where data was normalized, a Welch’s correction was used. For logarithmic data, values were first natural log (ln) transformed and then analyzed with T-tests. An ordinary one-way ANOVA with multiple comparisons was used to evaluate statistical differences among more than two groups with non-normalized data.

## ACKNOWLEDGMENTS

We would like to thank James Barber in the College of Veterinary Medicine flow cytometer core at the University of Georgia for his technical assistance. We also thank current and past members of the Brindley lab for helpful comments on the manuscript.

## AUTHOR CONTRIBUTIONS

Conceptualization, M.A.B., and J.M.R.B.; methodology, M.A.B., and J.M.R.B.; formal analysis, M.A.B. and J.M.R.B.; investigation, J.M.R.B., A.J.H., J.T.N., M.D.A., K.L.M., S.A.H., G.A.L.T., A.D., D.N.B., A.R.J., M.A.B.; data curation, J.M.R.B.; writing—original draft preparation, J.M.R.B.; writing—review and editing, J.M.R.B., A.J.H., J.T.N., M.D.A., K.L.M., S.A.H., G.A.L.T., A.D., D.N.B., A.R.J., M.A.B.; visualization, J.M.R.B., M.A.B.; supervision, M.A.B.; project administration, M.A.B.; funding acquisition, J.M.R.B., K.L.M., A.R.J., M.A.B. All authors have read and agreed to the published version of the manuscript.

---

**Supplemental Figure 1.**
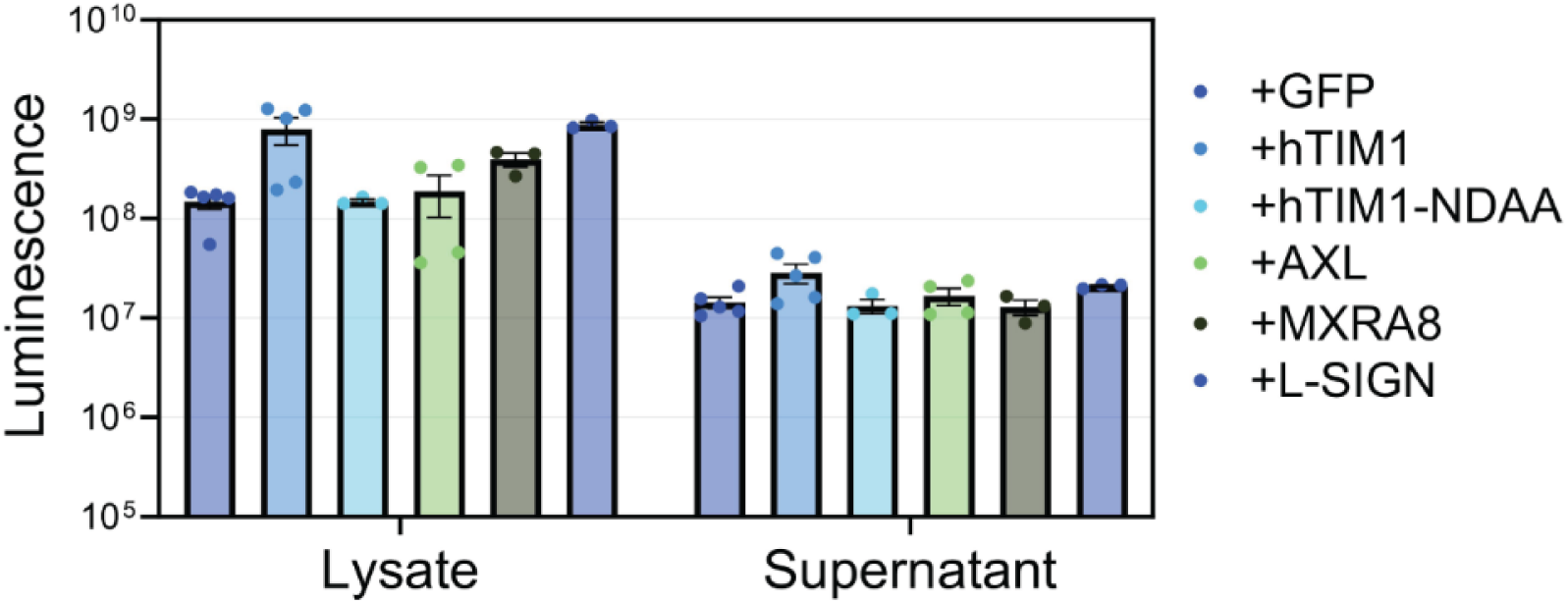
Exogenous expression of TIM, MXRA8, and L-SIGN increases cell lysate luminesce levels of CHIKV-infected cells. Correspo3nding levels of luminescence present in the total cell lysates and supernatants collected from the release efficiency assay shown in Figure 3E. Data represents the mean ±SEM from at least three independent trials.

**Supplemental Figure 2.**
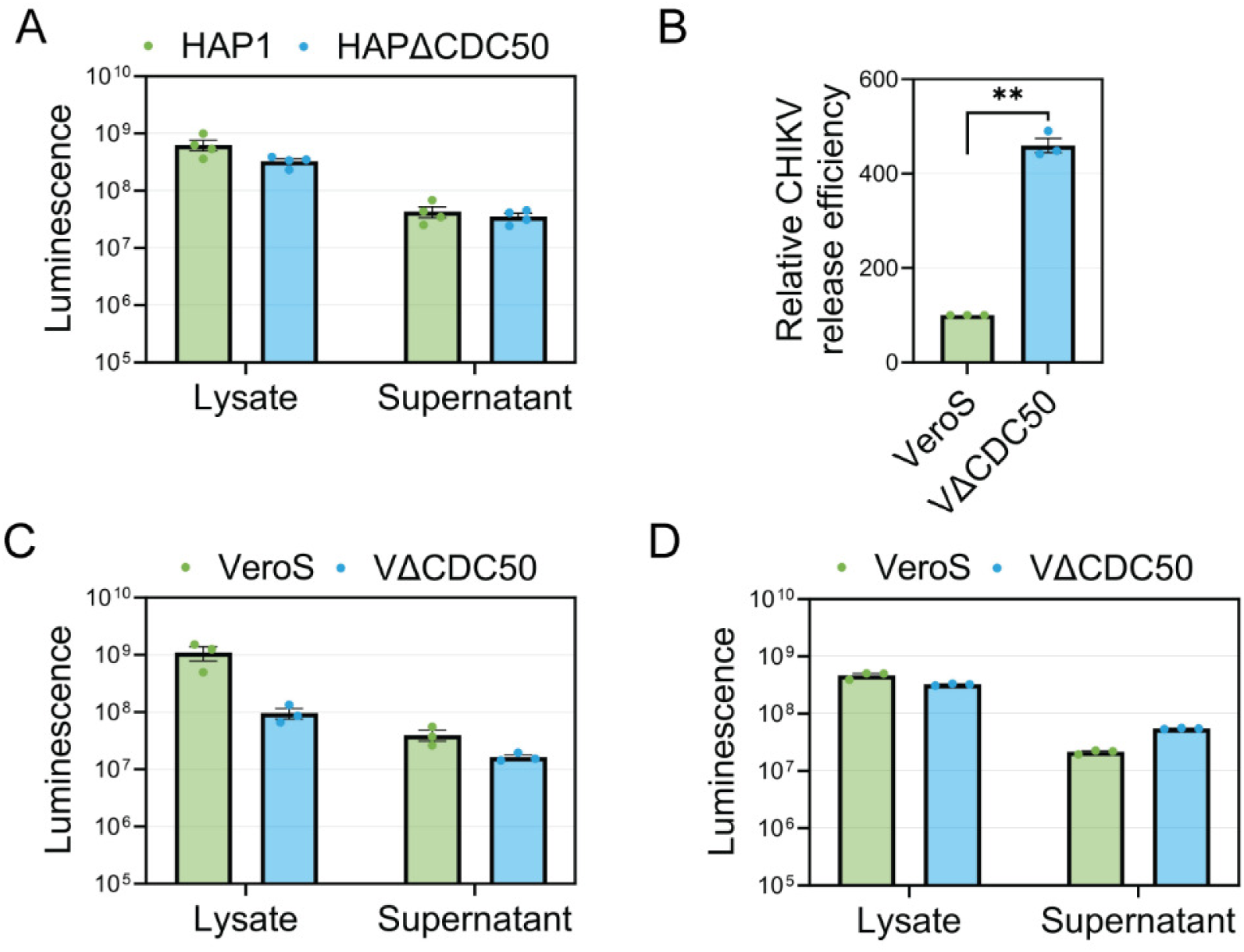
Raw luminescence levels of CHIKV-infected HAP1 and Vero cells knocked out for CDC50a. **(A)** Corresponding levels of luminescence present in the total cell lysates and supernatants collected from HAP1 and HAP1ΔCDC50 release efficiency assay shown in Figure 4C. **(B)** Corresponding levels of luminescence present in the total cell lysates and supernatants collected from VeroS and VeroSΔCDC50 release efficiency assay shown in Figure 5D or **(C)** with equalized cell lysate levels as shown in Figure 5E. Data represents the mean ±SEM from at least three independent trials.

**Supplemental Figure 3.**
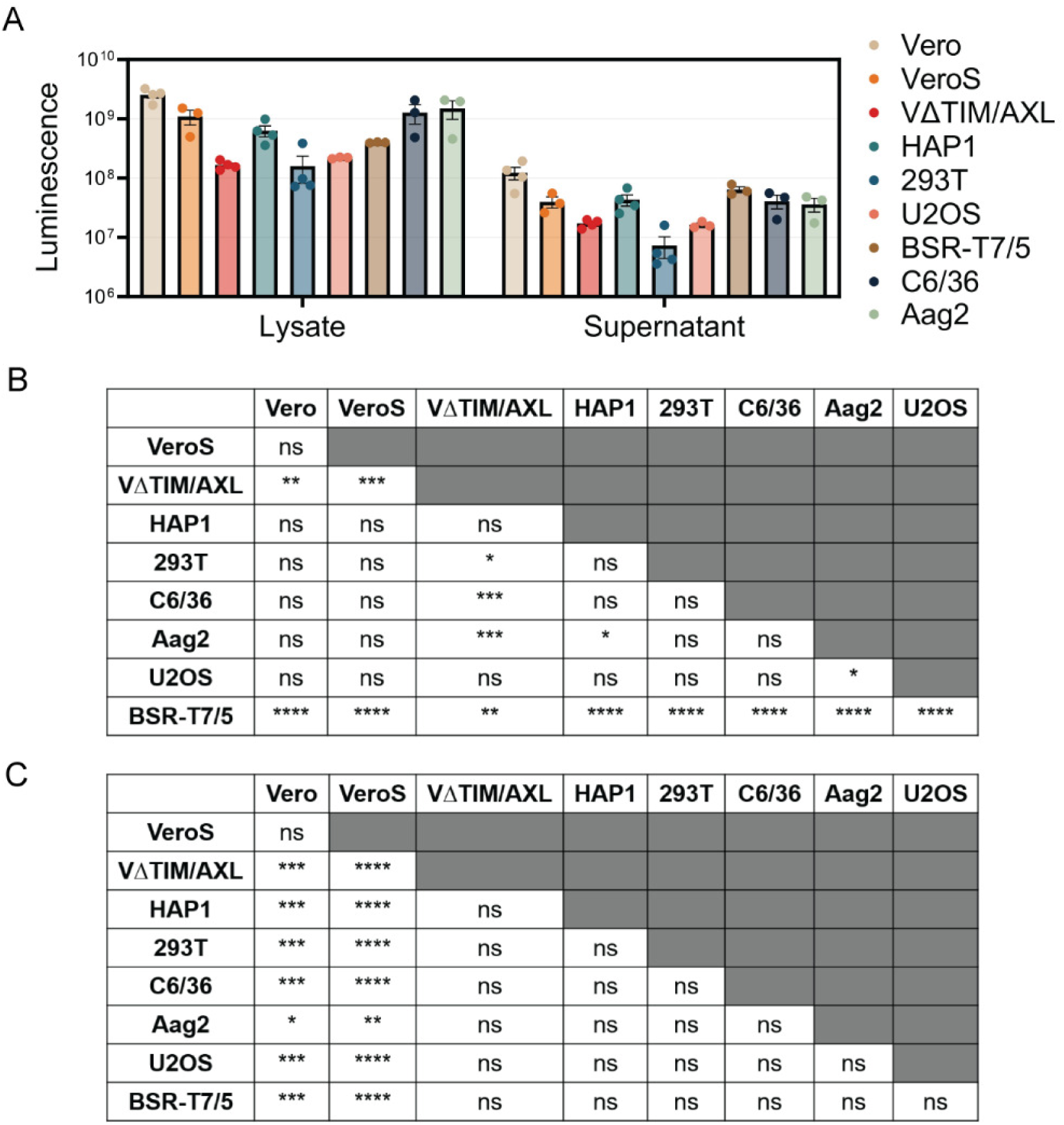
Luminescence levels from release efficiency assay in a panel of mammalian and mosquito cell lines. Corresponding levels of luminescence present in the total cell lysates and supernatants collected from release efficiency assay shown in Figure 6A. Data represents the mean ±SEM from at least three independent trials.

**Supplemental Figure 4.**
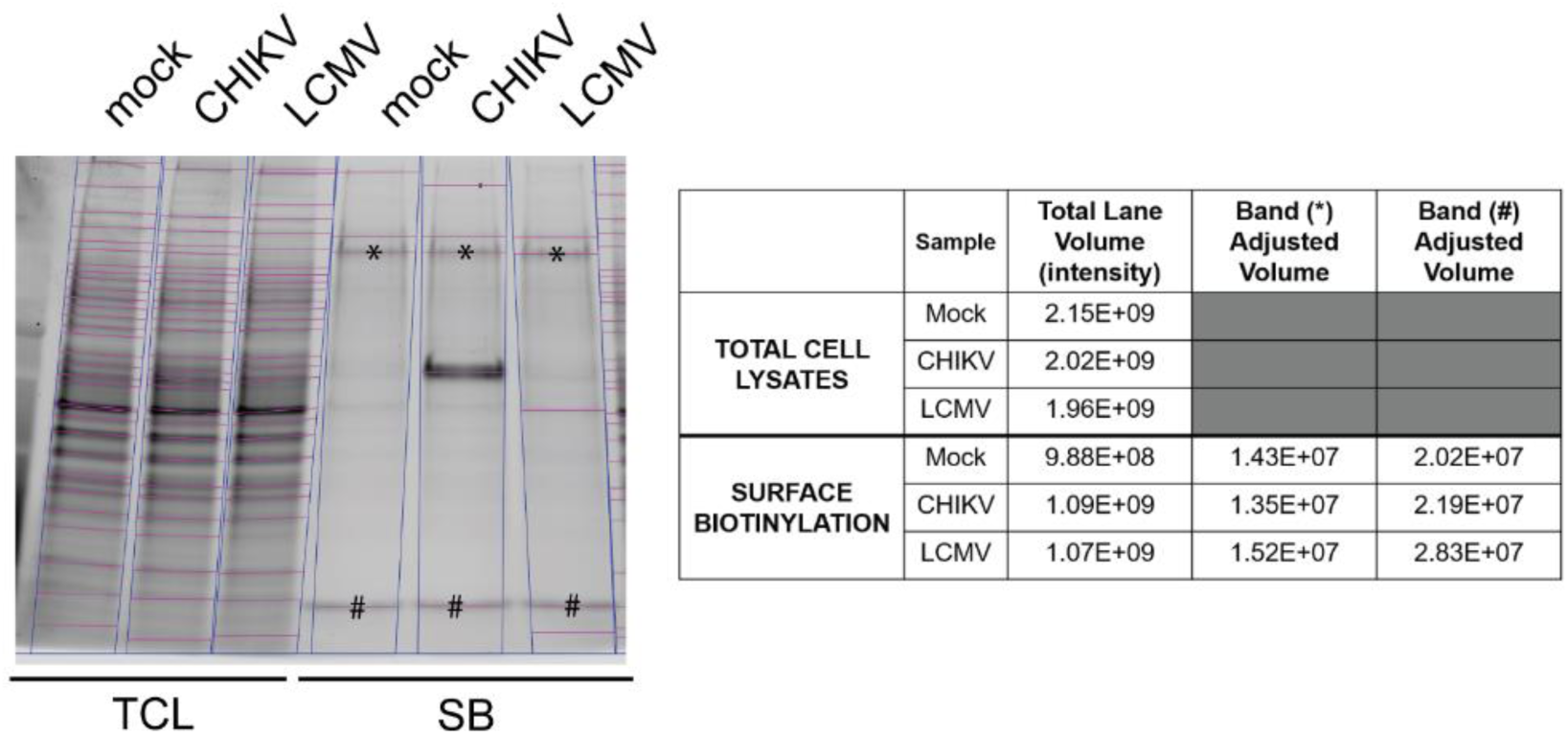
Quantification of surface biotinylation samples of CHIKV-infected cells. (A) Total cell lysate and surface biotinylated proteins of uninfected, CHIKV or LCMV-infected VeroS cells were quantified using densitometry analysis.

**Supplemental Figure 5.**
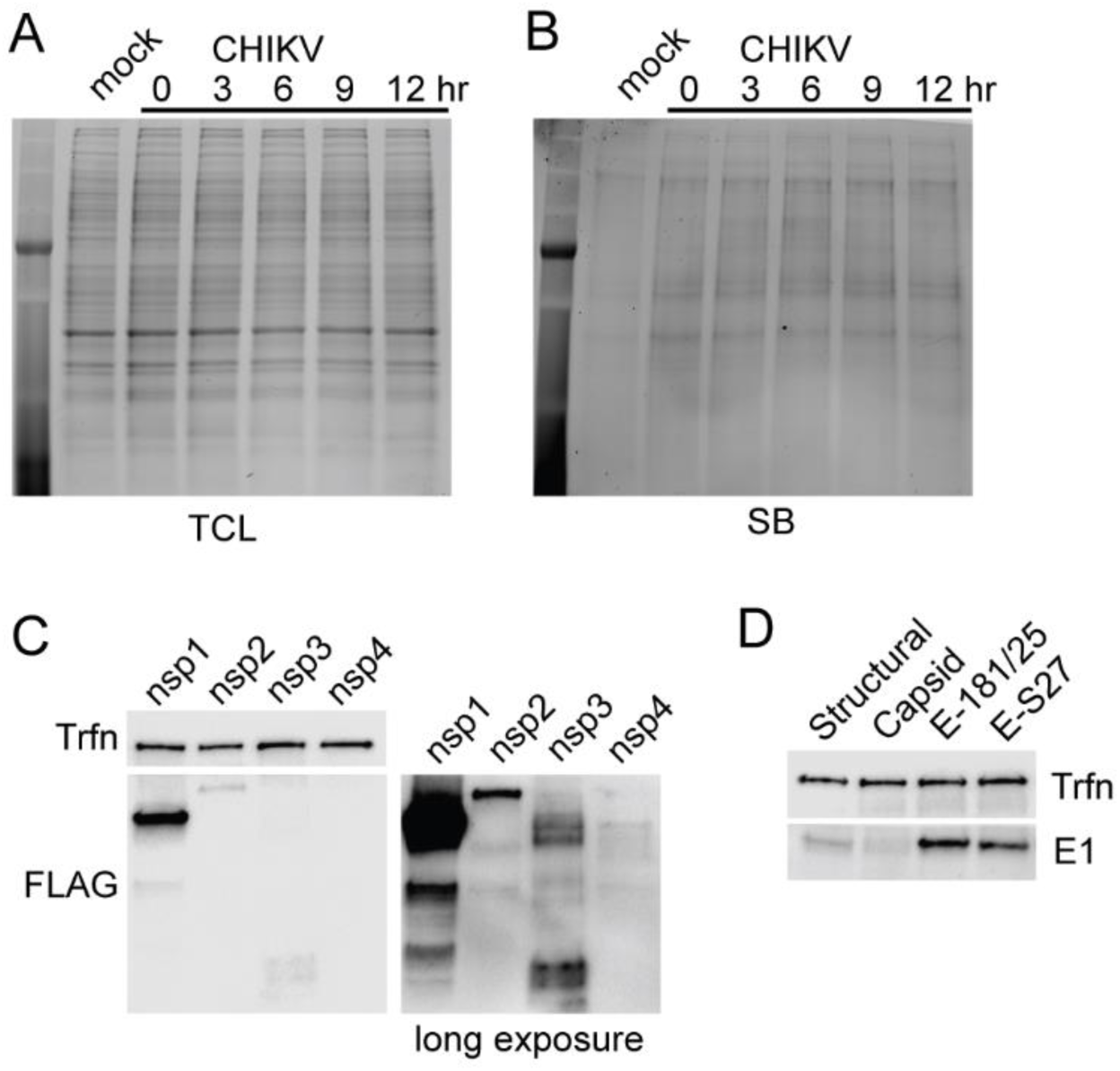
Production of exogenous expression of CHIKV proteins in VeroS cells. Stain-free gel analysis of **(A)** total cell lysates or **(B)** surface biotinylation proteins infected with CHIKV for different periods. **(C)** Exogenous expression of CHIKV non-structural proteins tagged with FLAG tag was analyzed through SDS-PAGE using an antibody against FLAG or transferrin as loading control. nsP1 was quickly detected in the cell lysates of transfection cells but longer exposure (right) was needed for detection of nsP2, nsP3, and nsP4. **(D)** Exogenous expression of CHIKV structural proteins was analyzed using an antibody against CHIKV E1 or transferrin as a loading control.

## References

1. Caglioti C, Lalle E, Castilletti C, Carletti F, Capobianchi MR, Bordi L. 2013. Chikungunya virus infection: an overview. New Microbiol 36:211–227.

2. Yap ML, Klose T, Urakami A, Hasan SS, Akahata W, Rossmann MG. 2017. Structural studies of Chikungunya virus maturation. Proc Natl Acad Sci 114:13703–13707.

3. Chmielewski D, Schmid MF, Simmons G, Jin J, Chiu W. 2022. Chikungunya virus assembly and budding visualized in situ using cryogenic electron tomography. Nat Microbiol 7:1270–1279.

4. Brown R, Wan J, Kielian M. 2018. The Alphavirus Exit Pathway: What We Know and What We Wish We Knew. Viruses 10:89.

5. Suomalainen M, Liljeström P, Garoff H. 1992. Spike protein-nucleocapsid interactions drive the budding of alphaviruses. J Virol 66:4737–4747.

6. Owen KE, Kuhn RJ. 1997. Alphavirus Budding Is Dependent on the Interaction between the Nucleocapsid and Hydrophobic Amino Acids on the Cytoplasmic Domain of the E2 Envelope Glycoprotein. Virology 230:187–196.

7. Ivanova L, Schlesinger MJ. 1993. Site-directed mutations in the Sindbis virus E2 glycoprotein identify palmitoylation sites and affect virus budding. J Virol 67:2546–2551.

8. Elmasri Z, Negi V, Kuhn RJ, Jose J. 2022. Requirement of a functional ion channel for Sindbis virus glycoprotein transport, CPV-II formation, and efficient virus budding. PLOS Pathog 18:e1010892.

9. Votteler J, Sundquist WI. 2013. Virus Budding and the ESCRT Pathway. Cell Host Microbe 14:232–241.

10. Jones PH, Maric M, Madison MN, Maury W, Roller RJ, Okeoma CM. 2013. BST-2/tetherin-mediated restriction of chikungunya (CHIKV) VLP budding is counteracted by CHIKV non-structural protein 1 (nsP1). Virology 438:37–49.

11. Ooi Y, Dubé M, Kielian M. 2015. BST2/Tetherin Inhibition of Alphavirus Exit. Viruses 7:2147–2167.

12. Kaletsky RL, Francica JR, Agrawal-Gamse C, Bates P. 2009. Tetherin-mediated restriction of filovirus budding is antagonized by the Ebola glycoprotein. Proc Natl Acad Sci 106:2886–2891.

13. Perez-Caballero D, Zang T, Ebrahimi A, McNatt MW, Gregory DA, Johnson MC, Bieniasz PD. 2009. Tetherin Inhibits HIV-1 Release by Directly Tethering Virions to Cells. Cell 139:499–511.

14. Acciani MD, Brindley MA. 2022. Scrambled or flipped: 5 facts about how cellular phosphatidylserine localization can mediate viral replication. PLoS Pathog 18:e1010352.

15. Leventis PA, Grinstein S. 2010. The Distribution and Function of Phosphatidylserine in Cellular Membranes. Annu Rev Biophys 39:407–427.

16. Segawa K, Nagata S. 2015. An Apoptotic ‘Eat Me’ Signal: Phosphatidylserine Exposure. Trends Cell Biol 25:639–650.

17. Reyes Ballista JM, Miazgowicz KL, Acciani MD, Jimenez AR, Belloli RS, Havranek KE, Brindley MA. 2023. Chikungunya virus entry and infectivity is primarily facilitated through cell line dependent attachment factors in mammalian and mosquito cells. Front Cell Dev Biol 11:1085913.

18. Moller-Tank S, Kondratowicz AS, Davey RA, Rennert PD, Maury W. 2013. Role of the phosphatidylserine receptor TIM-1 in enveloped-virus entry. J Virol 87:8327–41.

19. Kirui J, Abidine Y, Lenman A, Islam K, Gwon YD, Lasswitz L, Evander M, Bally M, Gerold G. 2021. The Phosphatidylserine Receptor TIM-1 Enhances Authentic Chikungunya Virus Cell Entry. Cells 10.

20. Li M, Ablan SD, Miao C, Zheng Y-M, Fuller MS, Rennert PD, Maury W, Johnson MC, Freed EO, Liu S-L. 2014. TIM-family proteins inhibit HIV-1 release. Proc Natl Acad Sci 111.

21. Xie S, Liang Z, Yang X, Pan J, Yu D, Li T, Cao R. 2021. Japanese Encephalitis Virus NS2B-3 Protein Complex Promotes Cell Apoptosis and Viral Particle Release by Down-Regulating the Expression of AXL. Virol Sin 36:1503–1519.

22. Ramjag A, Cutrone S, Lu K, Crasto C, Jin J, Bakkour S, Carrington CVF, Simmons G. 2022. A high-throughput screening assay to identify inhibitory antibodies targeting alphavirus release. Virol J 19:170.

23. McIntire JJ, Umetsu DT, DeKruyff RH. 2004. TIM-1, a novel allergy and asthma susceptibility gene. Springer Semin Immunopathol 25:335–348.

24. Moller-Tank S, Maury W. 2014. Phosphatidylserine receptors: Enhancers of enveloped virus entry and infection. Virology 468–470:565–580.

25. Zhang R, Kim AS, Fox JM, Nair S, Basore K, Klimstra WB, Rimkunas R, Fong RH, Lin H, Poddar S, Crowe JE, Doranz BJ, Fremont DH, Diamond MS. 2018. Mxra8 is a receptor for multiple arthritogenic alphaviruses. Nature 557:570–574.

26. Klimstra WB, Nangle EM, Smith MS, Yurochko AD, Ryman KD. 2003. DC-SIGN and L-SIGN Can Act as Attachment Receptors for Alphaviruses and Distinguish between Mosquito Cell- and Mammalian Cell-Derived Viruses. J Virol 77:12022–12032.

27. Shimojima M, Takada A, Ebihara H, Neumann G, Fujioka K, Irimura T, Jones S, Feldmann H, Kawaoka Y. 2006. Tyro3 family-mediated cell entry of Ebola and Marburg viruses. J Virol 80:10109–16.

28. Landi A, Iannucci V, Nuffel AV, Meuwissen P, Verhasselt B. 2011. One protein to rule them all: modulation of cell surface receptors and molecules by HIV Nef. Curr HIV Res 9:496–504.

29. Schneider-Schaulies J, Schnorr JJ, Brinckmann U, Dunster LM, Baczko K, Liebert UG, Schnei-der-Schaulies S, ter Meulen V. 1995. Receptor usage and differential downregulation of CD46 by measles virus wild-type and vaccine strains. Proc Natl Acad Sci U S A 92:3943–3947.

30. Breiner KM, Urban S, Glass B, Schaller H. 2001. Envelope protein-mediated down-regulation of hepatitis B virus receptor in infected hepatocytes. J Virol 75:143–150.

31. Li M, Waheed AA, Yu J, Zeng C, Chen H-Y, Zheng Y-M, Feizpour A, Reinhard BM, Gummuluru S, Lin S, Freed EO, Liu S-L. 2019. TIM-mediated inhibition of HIV-1 release is antagonized by Nef but potentiated by SERINC proteins. Proc Natl Acad Sci 116:5705–5714.

32. Stitt TN, Conn G, Gore M, Lai C, Bruno J, Radziejewski C, Mattsson K, Fisher J, Gies DR, Jones PF. 1995. The anticoagulation factor protein S and its relative, Gas6, are ligands for the Tyro 3/Axl family of receptor tyrosine kinases. Cell 80:661–670.

33. Morizono K, Xie Y, Olafsen T, Lee B, Dasgupta A, Wu AM, Chen ISY. 2011. The Soluble Serum Protein Gas6 Bridges Virion Envelope Phosphatidylserine to the TAM Receptor Tyrosine Kinase Axl to Mediate Viral Entry. Cell Host Microbe 9:286–298.

34. Bevers EM, Williamson PL. 2016. Getting to the Outer Leaflet: Physiology of Phosphatidylserine Exposure at the Plasma Membrane. Physiol Rev 96:605–645.

35. Kerr D, Gong Z, Suwatthee T, Luoma A, Roy S, Scarpaci R, Hwang HL, Henderson JM, Cao KD, Bu W, Lin B, Tietjen GT, Steck TL, Adams EJ, Lee KYC. 2021. How Tim proteins differentially exploit membrane features to attain robust target sensitivity. Biophys J 120:4891–4902.

36. Scott HL, Heberle FA, Katsaras J, Barrera FN. 2019. Phosphatidylserine Asymmetry Promotes the Membrane Insertion of a Transmembrane Helix. Biophys J 116:1495–1506.

37. Reitmayer CM, Levitt E, Basu S, Atkinson B, Fragkoudis R, Merits A, Lumley S, Larner W, Diaz AV, Rooney S, Thomas CJE, Von Wyschetzki K, Rausalu K, Alphey L. 2023. Mimicking superinfection exclusion disrupts alphavirus infection and transmission in the yellow fever mosquito Aedes aegypti. Proc Natl Acad Sci 120:e2303080120.

38. Cherkashchenko L, Rausalu K, Basu S, Alphey L, Merits A. 2022. Expression of Alphavirus Nonstructural Protein 2 (nsP2) in Mosquito Cells Inhibits Viral RNA Replication in Both a Protease Activity-Dependent and -Independent Manner. Viruses 14:1327.

39. Akhrymuk I, Lukash T, Frolov I, Frolova EI. 2019. Novel Mutations in nsP2 Abolish Chikungunya Virus-Induced Transcriptional Shutoff and Make the Virus Less Cytopathic without Affecting Its Replication Rates. J Virol 93:e02062–18.

40. Jose J, Taylor AB, Kuhn RJ. 2017. Spatial and Temporal Analysis of Alphavirus Replication and Assembly in Mammalian and Mosquito Cells. mBio 8:e02294–16.

41. Brouillette RB, Phillips EK, Patel R, Mahauad-Fernandez W, Moller-Tank S, Rogers KJ, Dillard JA, Cooney AL, Martinez-Sobrido L, Okeoma C, Maury W. 2018. TIM-1 Mediates Dystrogly-can-Independent Entry of Lassa Virus. J Virol 92.

42. Ono N, Tatsuo H, Hidaka Y, Aoki T, Minagawa H, Yanagi Y. 2001. Measles Viruses on Throat Swabs from Measles Patients Use Signaling Lymphocytic Activation Molecule (CDw150) but Not CD46 as a Cellular Receptor. J Virol 75:4399–4401.

43. Buchholz UJ, Finke S, Conzelmann K-K. 1999. Generation of Bovine Respiratory Syncytial Virus (BRSV) from cDNA: BRSV NS2 Is Not Essential for Virus Replication in Tissue Culture, and the Human RSV Leader Region Acts as a Functional BRSV Genome Promoter. J Virol 73:251–259.

44. Dorighello G, McPhee M, Halliday K, Dellaire G, Ridgway ND. 2023. Differential contributions of phosphotransferases CEPT1 and CHPT1 to phosphatidylcholine homeostasis and lipid droplet biogenesis. J Biol Chem 299:104578.

45. Lay Mendoza MF, Acciani MD, Levit CN, Santa Maria C, Brindley MA. 2020. Monitoring Viral Entry in Real-Time Using a Luciferase Recombinant Vesicular Stomatitis Virus Producing SARS-CoV-2, EBOV, LASV, CHIKV, and VSV Glycoproteins. Viruses 12.

46. Acciani MD, Lay Mendoza MF, Havranek KE, Duncan AM, Iyer H, Linn OL, Brindley MA. 2021. Ebola virus requires phosphatidylserine scrambling activity for efficient budding and optimal infectivity. J Virol JVI0116521.

47. Soh TK, Whelan SPJ. 2015. Tracking the Fate of Genetically Distinct Vesicular Stomatitis Virus Matrix Proteins Highlights the Role for Late Domains in Assembly. J Virol 89:11750–11760.

48. Cheng BYH, Ortiz-Riaño E, de la Torre JC, Martínez-Sobrido L. 2015. Arenavirus Genome Rearrangement for the Development of Live Attenuated Vaccines. J Virol 89:7373–7384.

49. Zhang L, Richard AS, Jackson CB, Ojha A, Choe H. 2020. Phosphatidylethanolamine and Phosphatidylserine Synergize To Enhance GAS6/AXL-Mediated Virus Infection and Efferocytosis. J Virol 95.

50. Salvador B, Zhou Y, Michault A, Muench MO, Simmons G. 2009. Characterization of Chikungunya pseudotyped viruses: Identification of refractory cell lines and demonstration of cellular tropism differences mediated by mutations in E1 glycoprotein. Virology 393:33–41.

